# Assessing risks and mechanisms of idiosyncratic drug toxicity by fingerprints of cell signaling responses

**DOI:** 10.1101/2022.12.01.518765

**Authors:** Alexander V. Medvedev, Sergei Makarov, Lyubov A. Medvedeva, Elena Martsen, Kristen L. Gorman, Benjamin Lin, Sergei S. Makarov

**Affiliations:** Attagene Inc., 7020 Kit Creek Rd, Morrisville NC 27560

## Abstract

Idiosyncratic drug-induced liver injury (DILI) is the leading cause of post-marketing drug withdrawal. Here, we describe a straightforward DILI liability assessment approach based on fingerprinting cell signaling responses. The readout is the activity of transcription factors (TF) that link signaling pathways to genes. Using a multiplex reporter assay for 45 TFs in hepatocytic cells, we assessed TF activity profiles (TFAP) for 13 pharmacological classes. The TFAP signatures were consistent with primary drug activity but transformed into different, ‘off-target’ signatures at certain concentrations (C_OFF_). We show that the off-target signatures pertained to DILI-relevant mechanisms, including mitochondria malfunction, proteotoxicity, and lipid peroxidation. Based on reported plasma concentrations in humans (C_MAX_), drugs do not reach the off-target thresholds in vivo, consistent with the lack of overt toxicity in the population. However, DILI liability drugs were dangerously close to the off-target thresholds. We characterized this closeness by the C_OFF_/C_MAX_ ratio termed the ‘safety margin’ (SM). Most-DILI-concern drugs invariably showed smaller safety margins than their less-concern counterparts in each pharmacological class and across classes (median SM values of 6.4 and 212.7, respectively (P<0.00015)). Therefore, the TFAP approach helps to explain idiosyncratic drug toxicity and provides clear quantitative metrics for its probability and the underlying mechanisms.

## INTRODUCTION

There are two kinds of drug toxicity(*1*). One is dose-dependent intrinsic toxicity, which occurs in most human patients and is usually detected in preclinical animal studies. The other is idiosyncratic drug toxicity (IDT), which occurs rarely and unpredictably, sometimes after several months of treatment(*2*), and is often detected after marketed drugs have gained broad exposure(*3*).

The IDT most frequently manifests as drug-induced liver injury (DILI), the leading cause of post-marketing drug withdrawal(*4*). DILI is a multi-factorial condition involving multiple mechanisms, such as mitochondrial damage, oxidative stress, endoplasmic reticulum stress, and disrupted bile acid homeostasis(*2, 5*). Furthermore, DILI is frequently accompanied by the immune response to drugs(*2*).

The most widely used DILI prediction approaches involve large panels of heterogeneous assays for multiple cell damages, e.g., to the mitochondrial function(*6*), reactive metabolite formation(*7*), immune and inflammatory biomarkers(*8*), and bile salt export pump (BSEP)(*9*). Furthermore, there are empiric rules (e.g., “rule of three”) predicting DILI liability by free radical formation, lipophilicity, and high drug concentration(*10*). However, the existing methods often fail to detect DILI-concern drugs, as evidenced by the post-marketing withdrawal of 462 drugs between 1953 and 2013(*4*)

Here, we describe an alternative evaluation approach based on fingerprinting cell signaling responses to drugs. These quantitative fingerprints characterize the activity of multiple transcription factors (TFs) connecting cellular signaling pathways to genes. The enabling technology is a multiplexed reporter assay (the FACTORIAL®) that permits quantitative parallel assessment of multiple TFs in a single well of cells(*11*).

We used the FACTORIAL platform in a previous study to survey TF responses to DILI-concern drug candidates(*12*). We found several preferentially responding to DILI-concern drugs TFs, but none of the individual TFs alone could unequivocally predict DILI liability.

This work focused on constellations of TF responses or TF activity profiles (TFAP) rather than individual TF responses. Using extensively validated FACTORIAL assay in the HepG2 hepatocyte cell line(*13, 14*), we examined TFAP signatures of 35 drugs from 13 pharmacological classes with diverse targets, chemistries, and MOA, categorized by the FDA LKTB database(*15*) as most-, less-, or no-DILI-concern drugs. The evaluated panel included withdrawn drugs and benchmark pairs of most-/less-DILI-concern drugs with identical MOA(*16*). Drugs were evaluated in a concentration-response format.

We found that each pharmacological class had a specific TFAP signature consistent with their primary activity. However, at some concentrations (C_OFF_), the ‘on-target’ signatures were transformed into different, ‘off-target’ signatures. These off-target signatures were shared by drugs of different classes, regardless of their pharmacology and DILI liability. To identify the underlying mechanisms, we queried previously described TFAP signatures of landmark perturbagens(*13*). The query showed that the most frequent off-target signatures were associated with DILI-relevant cell damage mechanisms, namely mitochondria malfunction, proteotoxicity, and lipid peroxidation.

Based on reported drug concentrations in plasma (C_MAX_), the bulk of evaluated drugs do not reach the C_OFF_ thresholds, consistent with the lack of overt adverse off-target effects in the population. However, the C_MAX_ values of most-DILI-concern drugs were much closer to the C_OFF_ thresholds than their safer counterparts. We characterized this proximity by the C_OFF_ /C_MAX_ ratio, which we termed the ‘safety margin’ (SM). Without exception, in each pharmacological class, most-DILI-concern drugs had smaller safety margins than less-concern drugs. Furthermore, across all drug classes, most-DILI-concern had significantly smaller median SM values than less-concern drugs (6.4 and 212.7, respectively (*P*<0.00015)). Therefore, the safety margin values reliably distinguish drugs with different DILI liabilities.

Therefore, the TFAP approach permits identifying underlying DILI mechanisms and provides clear quantitative metrics to aid in selecting and optimizing drug leads from various pharmacological classes.

## RESULTS

### Profiling TF responses by the FACTORIAL assay

The FACTORIAL is a multiplex reporter assay enabling quantitative parallel assessment of multiple TF activity in a single well of cells(*11*). The assay comprises TF-responsive constructs termed reporter transcription units (RTUs), identical to traditional reporter gene constructs, but we used reporter RNA transcripts as RTU activity readout. The FACTORIAL covers essential TF responses to xenobiotics, various stress stimuli, growth factors, and inflammation (listed in Fig. S1).

The assay workflow was described in previous publications(*11, 13*). It entails transient transfection of RTU plasmids into assay cells followed by parallel detection of RTU transcripts (Fig. S1). The homogeneous design of the FACTORIAL ensures equal detection efficacy across RTUs, drastically reducing the influence of experimental variables and resulting in exceptional reproducibility and the signal/noise ratio(*11*).

### Obtaining TFAP drug signatures

The evaluated panel comprised 35 drugs from 13 pharmacological classes categorized as most-, less-, and no-DILI-concern drugs by the FDA LTKB database(*15, 17*) (Table 1). Each evaluated class had less- and most-DILI-concern drugs with an identical mode of action (MOA).

**Table 1.** The DILI liability of evaluated drugs according to the FDA LKTB database(*15*).

| CLASS |  | DRUG | DILI concern |
| --- | --- | --- | --- |
| 1 | Glitazones | Troglitazone | Most (Withdrawn) |
|  |  | Rosiglitazone | Less |
|  |  | Pioglitazone | Less |
| 2 | Selective Estrogen Receptor modulators | Cyclofenil | Most (Withdrawn) |
|  |  | Tamoxifen | Most |
|  |  | Clomiphene | Less |
|  |  | Raloxifene | None |
| 3 | COMT inhibitors | Tolcapone | Most (Withdrawn) |
|  |  | Entacapone | None |
| 4 | Antibiotics (RNA-polymerase inhibitors) | Rifampicin | Most |
|  |  | Rifapentine | Most |
|  |  | Rifabutin | Less |
|  |  | Rifaximin | Less |
| 5 | Endothelin Receptor A antagonists | Sitaxentan | Most (Withdrawn) |
|  |  | Bosentan | Most |
|  |  | Ambrisentan | Less |
| 6 | Antibiotics (DNA gyrase inhibitors) | Trovafloxacin | Most (Withdrawn) |
|  |  | Moxifloxacin | None |
| 7 | Thrombin inhibitors | Ximelagatran | Most (Withdrawn) |
|  |  | Argatroban | None |
| 8 | Antiarrhythmic drugs | Benzbromarone | Most (Withdrawn) |
|  |  | Amiodarone | Most |
|  |  | Dronedarone | Most |
| 9 | Statins (HMG-CoA reductase inhibitors) | Atorvastatin | Most |
|  |  | Lovastatin | Less |
|  |  | Rosuvastatin | Less |
| 10 | 5-HT2A antagonists | Nefazodone | Most (Withdrawn) |
|  |  | Risperidone | Less |
| 11 | Vasodilators | Suloctidil | Most (Withdrawn) |
|  |  | Minoxidil | None |
| 12 | Anxiolytic | Alpidem | Most (Withdrawn) |
|  |  | Zolpidem | Less |
| 13 | SSRI | Zimelidine | Most (Withdrawn) |
|  |  | Fluvoxamine | Less |
|  |  | Fluoxetine | Less |

Drugs were assessed in a concentration-response format with serial dilutions, usually starting with a concentration of 20 to 60 µM until there were no discernable TF responses. If needed, we also assessed higher drug concentrations. As in previous studies(*12, 13*), we obtained the TFAP signatures after a prolonged (24 h) treatment to obviate differences in PKI pharmacokinetics and rapid transitory responses. Previously, we determined that this exposure time produced stable perturbagens’ signatures reflecting a steady activation state of cell signaling(*13*).

The PKI TFAP signatures are presented as radial graphs with log axes showing TF activity fold changes in PKI-treated vs. vehicle-treated cells. By this definition, the baseline TFAP signature in vehicle-treated cells is a perfect circle with R=1.0 (the “null” signature). Each presented here TFAP signature is an average of at least three independent FACTORIAL assays.

We assessed the pair-wise signatures’ similarity as the Pearson correlation coefficient r. We set the identity threshold at r=0.70 since the probability of two random 47-endpoint signatures correlating with r> 0.70 is very low (*P*<10-7) (*18*). Figs. 1-3 exemplify TFAP signatures of drugs from different pharmacological classes. For other drugs’ data, see Figs. S2-S12.

**Figure 1.**
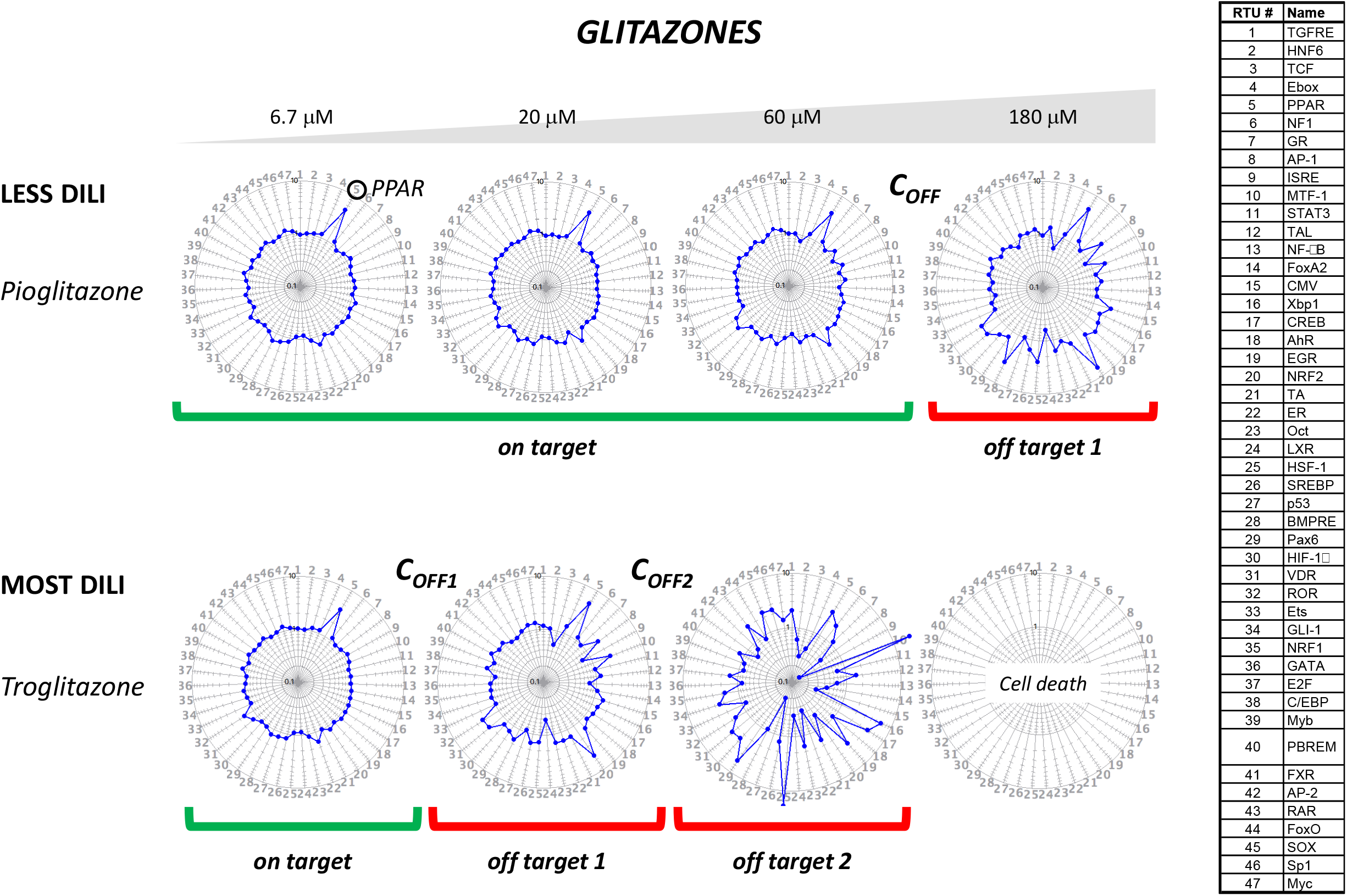
The TFAP signatures of glitazones. Drug signatures were obtained using the FACTORIAL assay in HepG2 cells after a 24-h treatment at indicated concentrations. The primary glitazone activity’s characteristic TF endpoint (PPAR) is circled in green. C_OFF_ indicates the on-target/off-target signature transition thresholds. Green and red lines indicate drug concentrations for the on-target and off-target signatures, respectively. Each TFAP signature is an average of at least three independent FACTORIAL assays.

**Figure 2.**
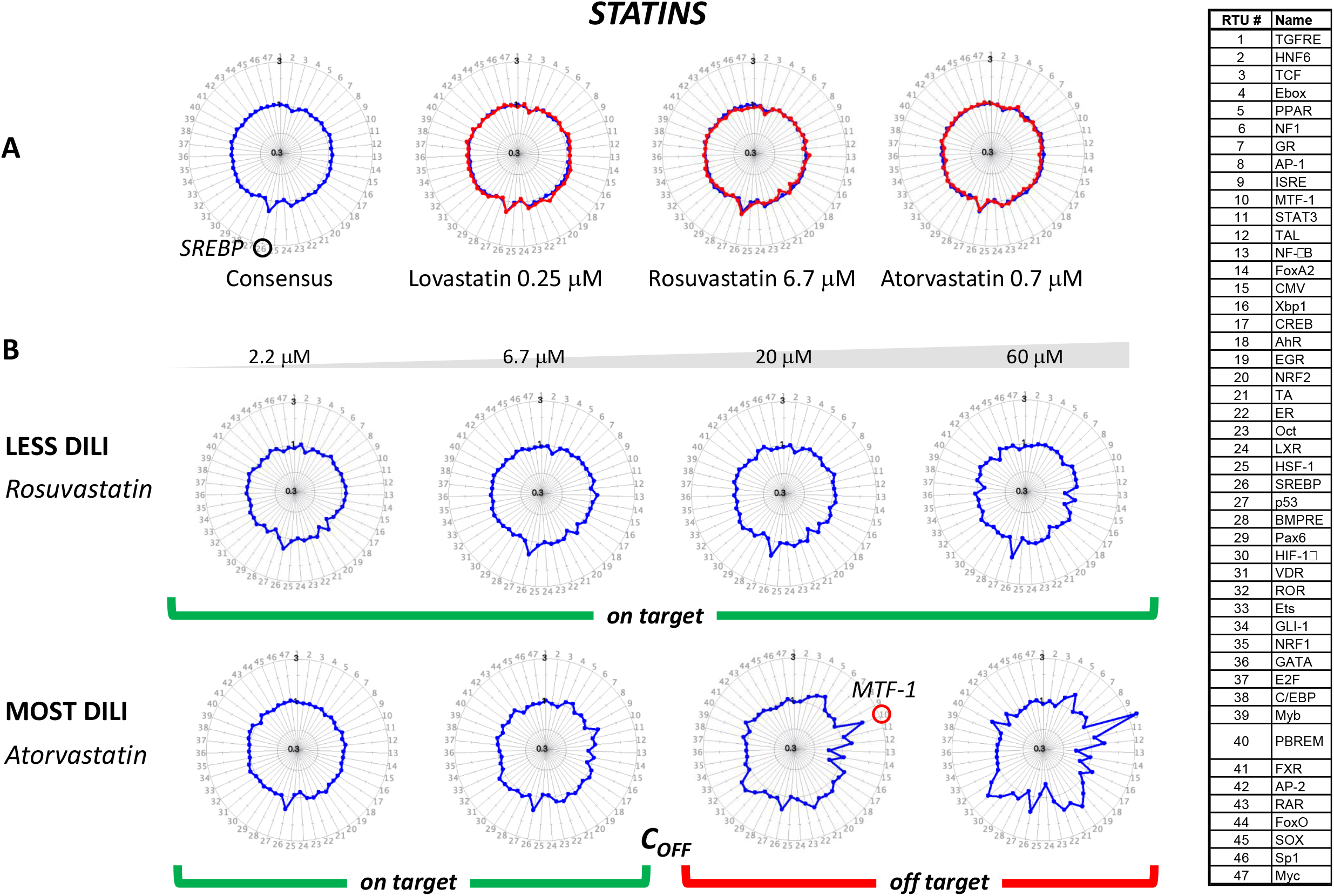
The TFAP signatures of statins. The drug signatures were obtained as described in Fig. 1 legend. **A**. The on-target signatures. The consensus signature of statins (blue) is overlayed by signatures of individual drugs (red). The characteristic endpoint of the primary activity (SREBP) is circled in green. **B**. Concentration-response signatures. C_OFF_ indicates the on-target/off-target signature transition threshold. Green and red lines indicate drug concentrations for the on-target and off-target signatures, respectively. The red circle indicates the off-target MTF-1 response. Each TFAP signature is an average of at least three independent FACTORIAL assays.

**Figure 3.**
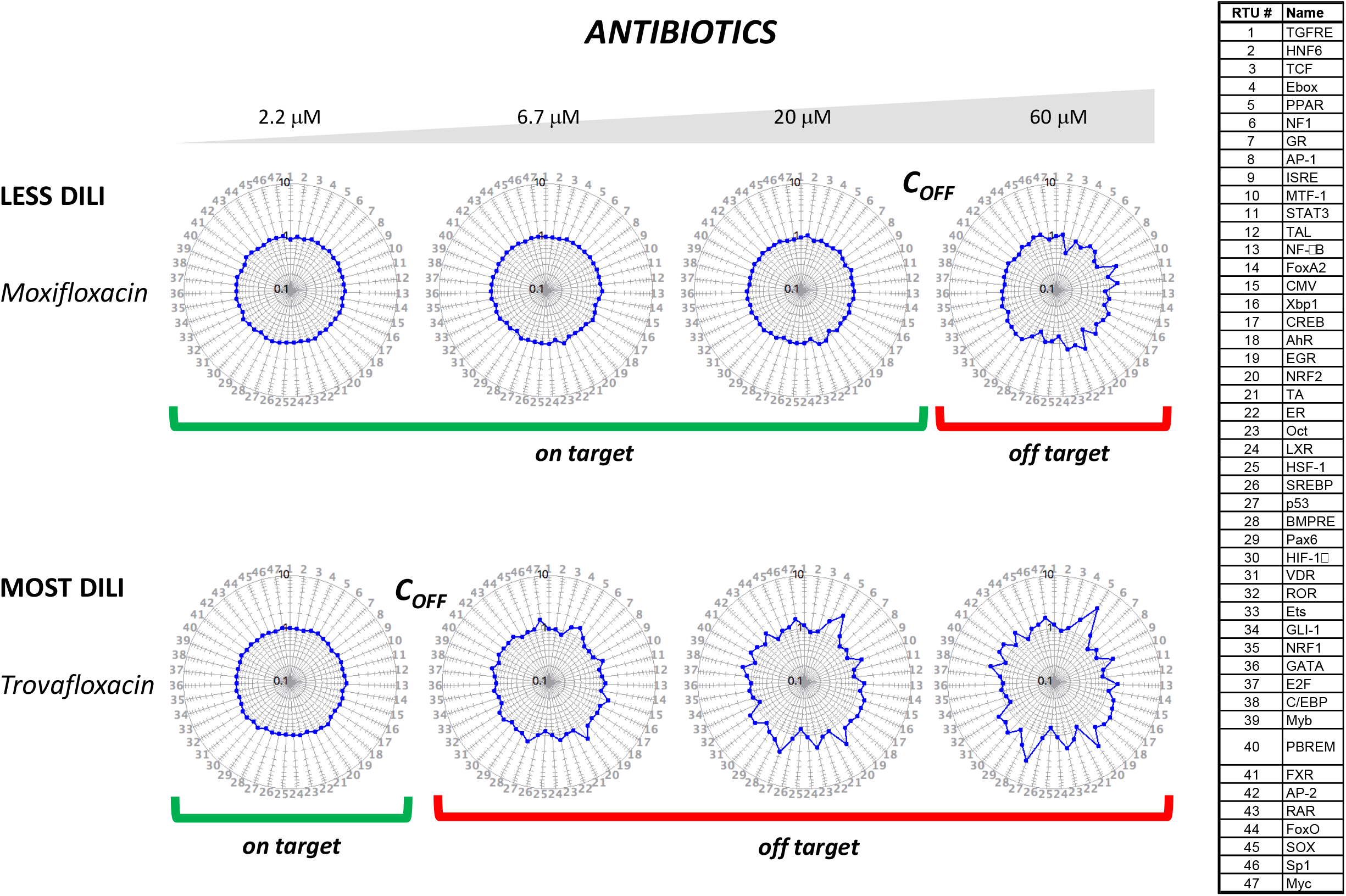
The TFAP signatures of fluoroquinolone antibiotics. The drug signatures were obtained as described in Fig. 1 legend. The C_OFF_ marks indicate the concentration threshold for the on-target (null signature) to off-target signature transition. Green and red lines indicate drug concentrations for the on-target and off-target signatures, respectively. Each TFAP signature is an average of at least three independent FACTORIAL assays.

#### Glitazones

Glitazones are agonists of peroxisome proliferator-activated receptor gamma (PPARg) used for treating type II diabetes(*19*). Therefore, the PPAR RTU of the FACTORIAL provides a direct readout for their primary activity. We assessed concentration-response signatures of troglitazone (most-DILI concern, withdrawn due to hepatotoxicity) and less-DILI concern pioglitazone and rosiglitazone. Fig. 1 shows the signatures of pioglitazone and troglitazone (for rosiglitazone signature, see Fig. S2A). At lower concentrations, drugs exhibited identical signatures consistent with their primary activity at PPAR (RTU#5). However, after reaching certain thresholds (C_OFF_), drugs produced different, ‘off-target’ signatures with additional TF responses, including activated oxidative stress-responsive NRF2/ARE and bone morphogenic protein response element (BMPRE) RTUs (# 20 and 28). With further concentration increase, troglitazone exhibited a secondary off-target signature (off-target TFAP2) (Fig. 1).

All glitazones had identical on-target and off-target TFAP signatures, differing only in the signature transition values (C_OFF_ ∼20 µM for troglitazone and ∼180 µM for pioglitazone and rosiglitazone). According to literature, maximal plasma concentrations (C_MAX_) of rosiglitazone, pioglitazone, and troglitazone in humans (1.0 µM, 2.9 µM, and 6.4 µM, respectively (Table S1)) are below the C_OFF_ thresholds. However, the C_MAX_ of troglitazone was much closer to the C_OFF_ threshold vs. less-DLI-concern pioglitazone and rosiglitazone.

#### Statins

Widely prescribed statins account for about 7% of all registered DILI incidents(*20*). Here, we compared TFAP signatures of atorvastatin (most-DILI concern) and less-DILI concern rosuvastatin and lovastatin.

The cholesterol-lowering activity of statins is due to inhibited cholesterol synthesis in the liver. The cholesterol reduction activates a feedback mechanism that causes proteolytic activation of TFs called sterol regulatory element-binding proteins (SREBP)(*21*). Thus, the SREBP RTU activity provides a direct readout for statins’ primary activity. We found that all evaluated statins at low concentrations shared a consensus TFAP signature reflecting the on-target activity at SREBP (Fig. 2A).

The concentration-response analysis revealed differences in cell responses to statins. Rosuvastatin exhibited the on-target signature at all tested concentrations, up to 60 µM. In contrast, atorvastatin elicited a distinct off-target signature at 20 µM and lovastatin at 7.4 µM (Fig. S2). The most pronounced feature of atorvastatin off-target signature was activation of metal-responsive transcription factor 1 (MTF-1) (RTU#10), which was consistent with the report showing induction of an MTF-1 target gene metallothionein by high atorvastatin doses in mice(*22*).

Based on reported plasma concentrations (C_MAX_) in humans, therapeutic concentrations of rosuvastatin (0.01 µM), lovastatin (0.01 µM), and atorvastatin (0.12 µM) (Table S1) do not reach the off-target thresholds. Nonetheless, the C_MAX_ value for most-DILI-concern atorvastatin was closer to the C_OFF_ threshold than less-concern rosuvastatin and lovastatin.

#### Antibiotics

Trovafloxacin (most-DILI-concern, withdrawn) and moxifloxacin (no-DILI concern) are inhibitors of bacterial DNA gyrase and topoisomerase IV(*23*). As human cells lack these targets, the expected on-target TFAP signature of these drugs was the null signature.

Both drugs showed the null signature until certain concentration thresholds (Fig. 3). At 6.7 µM, trovafloxacin elicited multiple TF responses, including activation of PPAR (#5), NRF2 (#20), Oct (#23), and p53 RTUs (#27). By comparison, moxifloxacin elicited detectable TF responses only at 60 µM, and these responses differed from those by trovafloxacin (Fig. 3). The reported C_MAX_ value for both drugs is ∼7 µM (Table S1), suggesting that trovafloxacin can reach the C_OFF_ threshold in vivo.

### The off-target TFAP signatures epitomize cell responses to DILI-relevant damage mechanisms

Our data (Figs. 1-3 and S2-12) show that the signatures of drugs from different pharmacological classes have similar concentration-response patterns. Namely, until certain concentrations, these signatures were consistent with the primary drug activity but then transformed into non-specific off-target signatures. To reveal the underlying off-target signatures mechanisms, we queried our reference signature dataset of landmark perturbagens.

This dataset comprises TFAP signatures of various chemicals and landmark perturbagens, most of them evaluated under the ToxCast project(*24*). This dataset comprises multiple signature clusters, and some annotated cluster signatures were described in our previous publication, including consensus TFAP signatures for mitochondria and HDAC inhibitors, Ub/proteasome inhibitors, cytoskeleton disruptors, and DNA damaging agents(*13*). Importantly, these signatures permitted straightforward identification of compounds with given bioactivity among uncharacterized chemicals(*13*). Thus, we queried the off-target drug signatures of this study against the reference landmark signatures to ascertain the underlying mechanisms.

#### Mitochondria malfunction

Previously, we described the consensus signature shared by different inhibitors of the mitochondrial electron transfer chain(*13*). This signature epitomizes known retrograde signaling mechanism of adaptive cell response to mitochondria damage(*25*). Most drugs of this study (21 out of 35) at certain concentrations had identical signatures (r>0.70) to the mitochondria damage response signature (Fig. 4). Notably, the inferred mitochondria inhibiting activity agreed with the literature data on these drugs (Fig. 4).

**Figure 4.**
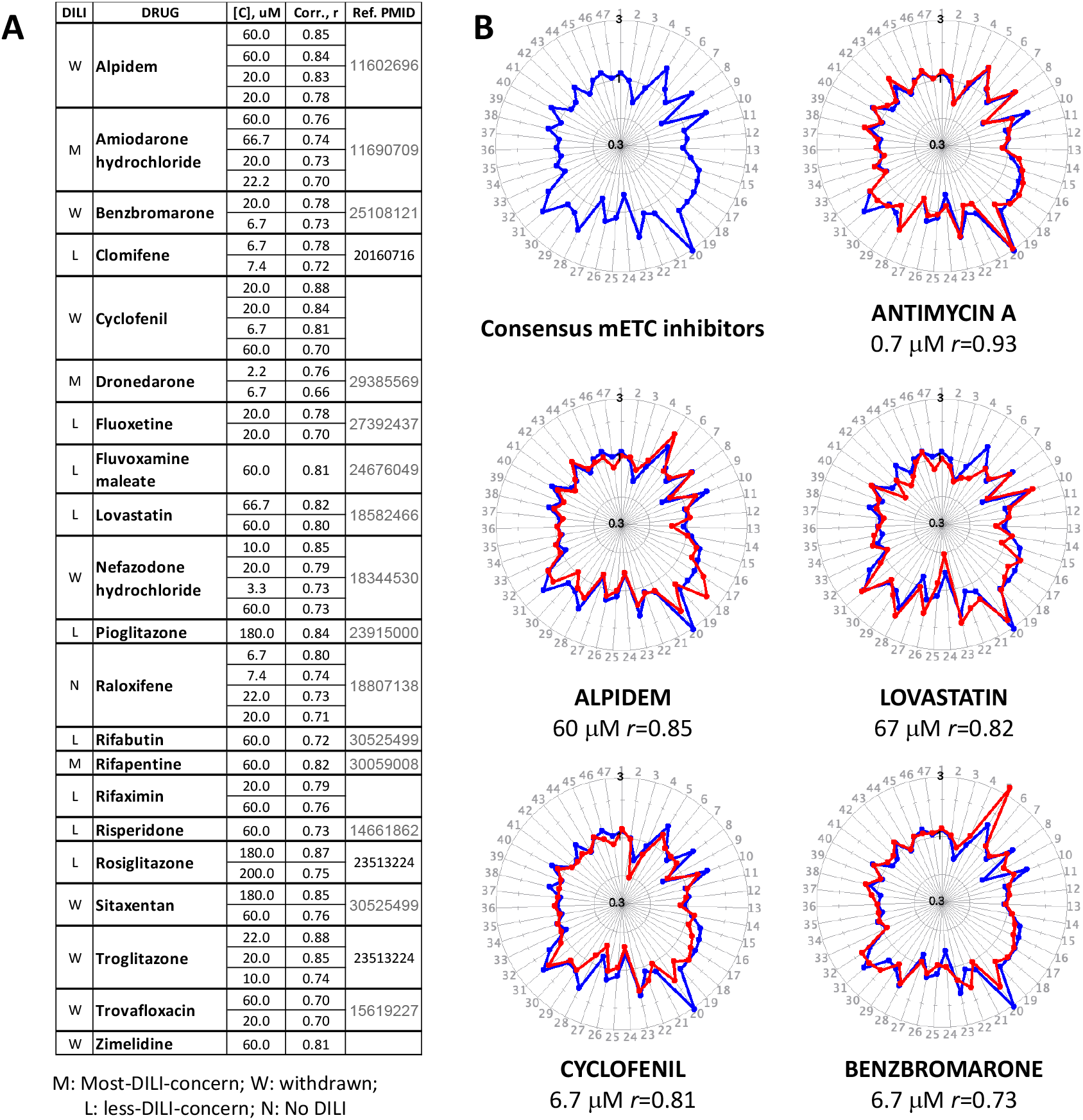
Drugs with the mitochondria damage response signature. Drug signatures were queried against the consensus signature of mitochondria inhibitors(*13*). **A**. The list of retrieved drugs with identical (r>0.70) signatures. The similarity values at a given drug concentration are shown. Identical concentrations relate to data from independent experiments. The Ref. PMID column lists the PMID # of the literature references for the mitochondria-inhibiting activity of corresponding drugs. **B**. The consensus signature of mitochondria inhibitors (blue) overlaid with the signature of a prototypical mitochondria inhibitor Antimycin A and indicated drugs (red). The similarity values for the overlaid signatures at indicated concentrations are shown. Each TFAP signature is an average of at least three independent FACTORIAL assays.

#### Proteotoxicity

Our previous publication also described the consensus TFAP signature for inhibitors of the ubiquitin-proteasome (Ub/PS) protein degradation pathway(*13*). Subsequently, we found other landmark perturbagens with this signature that did not inhibit the Ub/PS pathway, including cadmium, hydrogen peroxide, and geldanamycin (Fig. 5A,B). The unifying feature of these compounds and Ub/PS inhibitors is that they all disrupt protein homeostasis (proteostasis) through accumulation of misfolded proteins. For example, Ub/PS inhibitors, such as bortezomib, prevent misfolded protein degradation(*26*); geldanamycin inhibits Hsp90 chaperone required for protein folding(*27*); cadmium ions interfere with protein folding during translation(*28*), and hydrogen peroxide causes protein oxidation(*29*). Therefore, this consensus signature represents a bona fide marker of cell response to compound proteotoxicity. The proteotoxicity response signature was found ten drugs of this study with different DILI liabilities (Fig. 5C).

**Figure 5.**
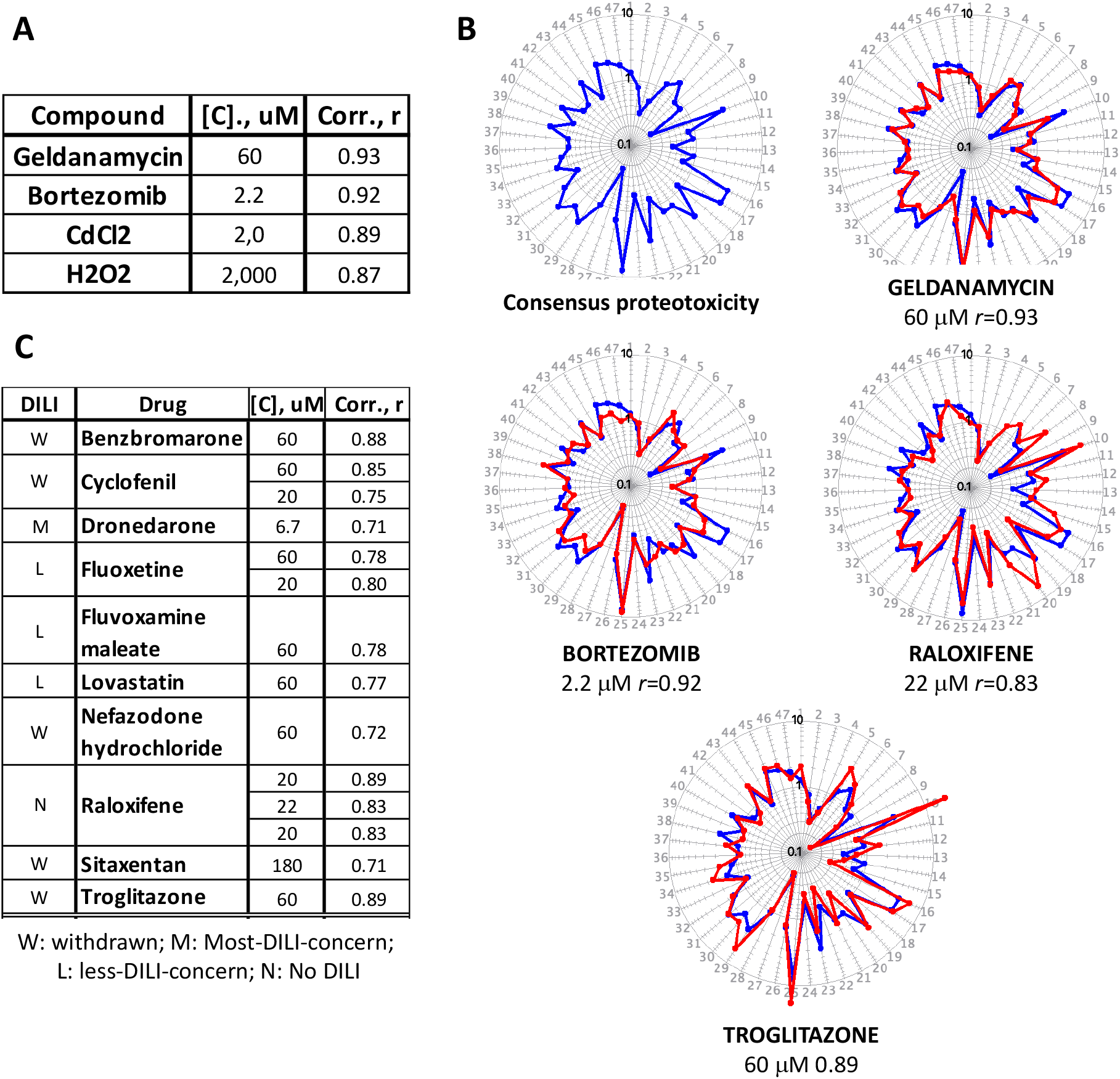
Drugs with the proteotoxicity response signature. **A**,**B.**The signature of cell response to proteotoxic compounds, a.k.a. the consensus signature of Ub/proteasome inhibitors(*13*). **C**. The list of retrieved drugs with identical (r>0.70) signatures. The consensus signature (blue) is overlaid by individual signatures (red). The r values are Pearson correlation of the overlaid signatures. **D**. Drugs with the signature of mitochondria damage response exhibit the proteotoxicity signature at higher concentrations. The correlation values *r* with indicated consensus signatures are shown. Each TFAP signature is an average of at least three independent FACTORIAL assays.

#### Lipid peroxidation

One signature cluster found in our reference dataset was tentatively annotated as the cluster of lipid peroxidation (LPO) inducers. This annotation was based on the presence of two lipophilic radical initiators 2,2′-azobis(2,4-dimethylvaleronitrile) (AMVN) and 2,2’-azodi(2-methylbutyronitrile) (AMBN) (Fig. 6A,B). These unstable compounds decompose in water solutions at constant rates, producing hydrophobic free radicals(*30, 31*). These radical initiators are used as specific inducers for studying lipid peroxidation in biological membranes(*30, 31*). The consensus LPO signature (Fig. 6B) was found for six drugs of this study with different DILI liabilities (Fig. 6A,B). Notably, all these drugs are highly lipophilic chemicals (logP>4), further supporting the relevance of the consensus LPO signature to membrane damage response.

**Figure 6.**
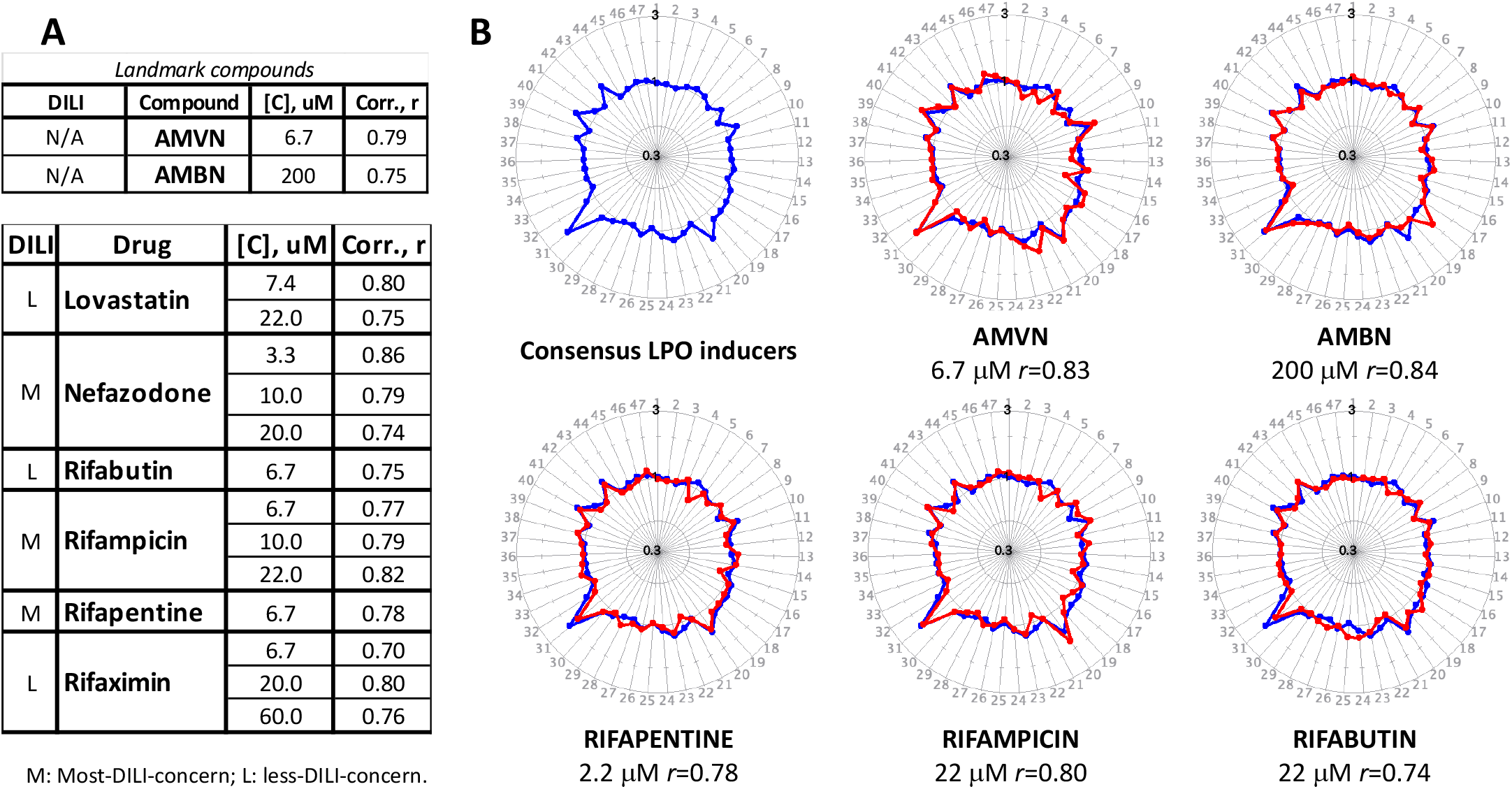
Drugs with the lipid peroxidation response signature. **A.** The similarity values of the putative consensus signature of lipid peroxidation (LPO) response to the signatures of landmark LPO inducers (upper panel) and drugs of this study (bottom) panel.). **B**. The consensus signature (blue) is overlaid with the signatures of landmark LPO inducers and evaluated drugs (red). **C**. Drugs with the signature of LPO response exhibit the signature of mitochondria damage response at higher concentrations. Each TFAP signature is an average of at least three independent FACTORIAL assays.

### The off-target signatures indicate a hierarchy of responses to DILI damage mechanisms

Concentration-response analysis showed that drug signatures at different concentrations reflected cell responses to different damage mechanisms. For example, nefazodone signatures matched the LPO signature at a lower concentration and the mitochondria damage response signature at a higher concentration (Fig. 7A). We observed this off-target signature transition in 5 out of 6 drugs with the LPO signature (compare drug lists of Figs. 6 and 4). In each case, the LPO signature preceded the signature of mitochondria damage response.

Furthermore, many drugs with the signature of mitochondria damage response showed at higher concentrations the proteotoxicity response signatures. For example, see the off-target signatures of troglitazone (Fig. 7B). In addition, some drugs at certain concentrations exhibited transitory signatures that matched both the consensus signatures of mitochondria damage and proteotoxicity response (compare the lists of drugs in Figs 4A and 5A).

**Figure 7.**
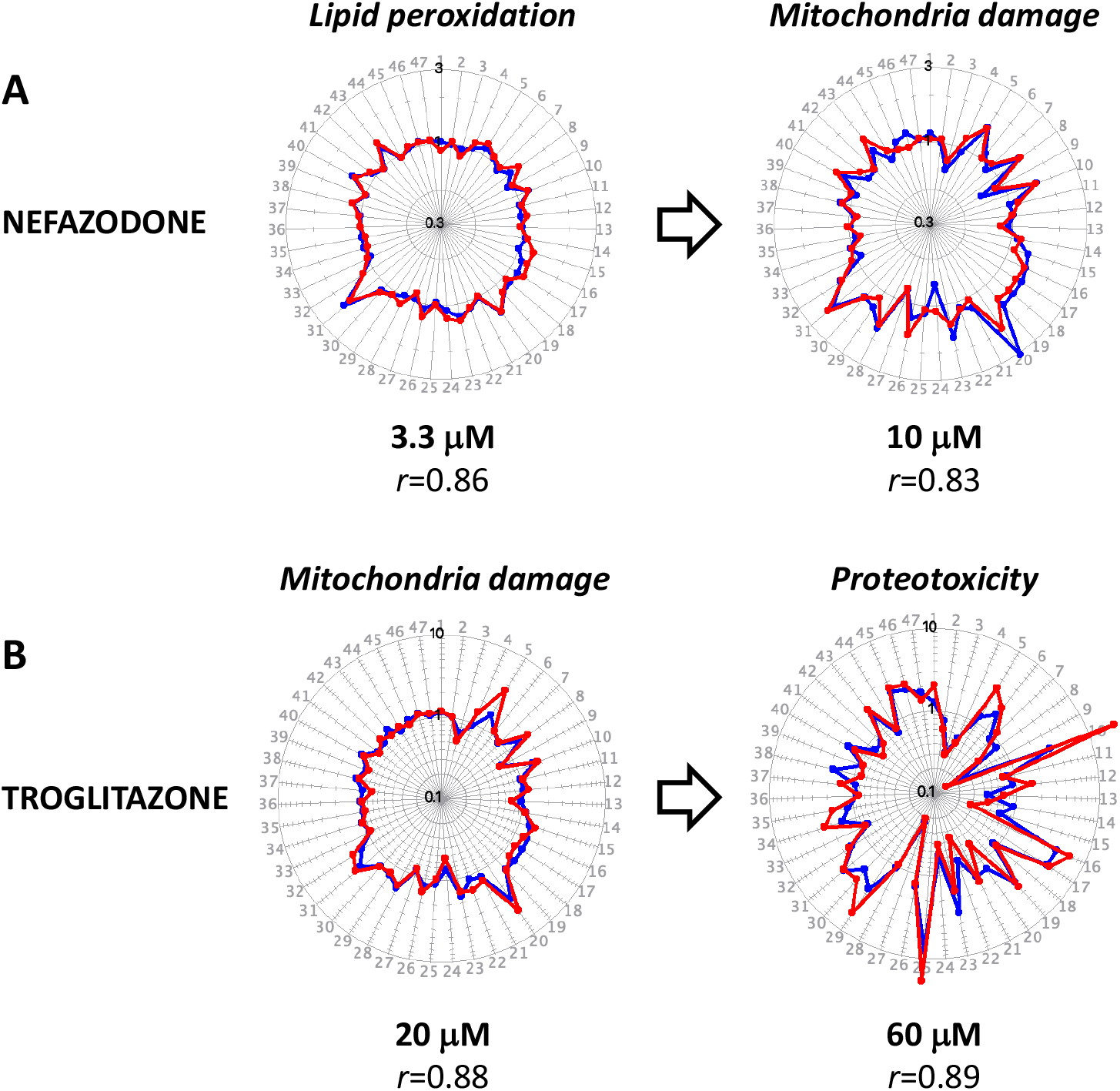
Drug safety margin as a quantitative descriptor of DILI liability. **A**. The evaluated drugs’ safety margin (SM) values (C_OFF_/C_MAX_ ratio). **B**. The SM values for drugs with different DILI liabilities across the pharmacological classes. The dotted line indicates the SM value of 1.0. Note that most-DILI-concern drugs of each class have lower safety margins than their safer counterparts, with the lowest SM values for withdrawn drugs. **C**. The median SM values for the withdrawn/most DILI and less/no DILI drugs across classes. The statistical significance is calculated by a two-sided Mann-Whitney test.

These observations align with known mechanisms of DILI pathology. Lipid peroxidation can lead to mitochondria malfunction(*32, 33*), and mitochondrial stress can cause protein unfolding(*34, 35*). Furthermore, the off-target signature transition implies a hierarchy of cell damage responses. The LPO signature indicates cell response to drug-induced membrane damage, but the resulting mitochondria damage switches cells’ response to accommodate the mitochondrial stress. The dominant cell response to mitochondria malfunction-induced protein damage accommodates the disrupted proteostasis.

Therefore, the off-target drug signatures signify DILI-relevant cell damage responses. However, as drugs with different DILI liabilities exhibited these signatures, we searched for a specific DILI liability marker.

### DILI liability correlates with the closeness of circulating drug concentrations to the off-target thresholds

Based on literature analysis, reported drug concentrations in plasma (C_MAX_) (Table S1) for most evaluated drugs were below the off-target effect thresholds in assay cells (C_OFF_). That was consistent with the lack of conspicuous off-target adverse drug reactions in the large population. However, the C_MAX_ values of most-DILI-concern drugs were closer to C_OFF_ values than less-concern drugs. We characterized this proximity by the C_OFF_ /C_MAX_ ratio that we termed ‘the safety margin,’ or SM. The SM value of 1.0 separates drugs that do not reach the off-target threshold (SM>1.0) from drugs that do (SM<1.0).

Only two of 35 evaluated drugs had SM values<1.0, both of them most-DILI-concern drugs (Figs. 7A-B). Importantly, in each pharmacological class, most-DILI-concern drugs invariably had smaller safety margins than their less- and no-DILI-concern counterparts, with smallest SM values for withdrawn drugs (Fig. 7B). Across drug classes, the median safety margin of most-DILI-concern drugs was more than 30-fold smaller than less- and no-DILI-concern drugs (6.4 vs. 212.7, respectively (P<0.00015)) (Fig. 7C). Therefore, the safety margin provides a clear-cut quantitative parameter distinguishing most- and less-DILI-concern drugs.

## DISCUSSION

Despite the significant progress in understanding DILI pathology, assessing DILI liability at early drug development stages remains an acute problem. Here, we presented a straightforward DILI liability evaluation approach based on fingerprinting drug-induced cell signaling responses. To capture these responses, we used a multiplex reporter system (the FACTORIAL) for 45 TFs. Using this system in human hepatocyte HepG2 cells, we assessed TF activity profiles (TFAP) for 35 drugs of 13 pharmacological classes. The main results are as follows.

1. The TFAPs provided quantitative and reproducible descriptors of cell response to drugs. Within certain concentration ranges, drugs showed specific TFAP signatures consistent with their primary activity. For example, glitazones’ signatures reflected the on-target activity at PPAR (Fig. 1 and S2), and statins’ signatures reflected inhibited cholesterol synthesis epitomized by SREBP activation (Figs. 2 and S2). When the assay cells did not express drug targets (e.g., antibiotics), the on-target signature was the null signature (Figs. 3 and S5). Furthermore, we showed elsewhere that this approach applies to other drug classes with specific on-target TFAP signatures, such as kinase inhibitors(*36*).
2. After reaching certain concentration thresholds (C_OFF_), drugs elicited non-specific ‘off-target’ signatures shared by drugs of different classes. To ascertain their biological meaning, we queried reference TFAP signatures of landmark perturbagens. This analysis revealed three main off-target signatures. The most frequent off-target signature, shared by ∼ 61% of drugs, signified mitochondria malfunction. Notably, the inferred mitochondria inhibiting activity agreed with the reported data by others (Fig. 4). Two other off-target signatures, exhibited by ∼ 29% and ∼ 17% evaluated drugs, respectively, signified cell responses to proteotoxicity (Fig. 5) and lipid peroxidation (Fig. 6). The off-target signatures were consistent with known free-radical mechanisms of DILI pathology. Mitochondria damage is the hallmark of DILI(*2, 3, 37, 38*). Disrupted proteostasis, manifested by protein unfolding, is another characteristic feature of DILI(*39, 40*). Because of that, reactive metabolite formation is an essential criterium for selecting drug leads(*7*). Lipid peroxidation, too, has been associated with liver injury by lipophilic radicals(*41, 42*) that trigger lipid peroxidation in cell membranes, causing increased membrane permeability and disrupting membrane-associated proteins(*43*). In line with that, high drug lipophilicity correlates with DILI liability(*44*). As described previously, perturbagens of many biological processes and cell systems have specific TFAP signatures that epitomize coordinated signaling responses to accommodate cell damages(*13*). Data of this study indicate a hierarchy of these responses to drug toxicity. For example, most drugs with the LPO signature at higher concentrations exhibited the signature of mitochondria damage response (Fig. 7A). Akin to that, the signature of mitochondria damage response at higher concentrations often transformed into the signature of proteotoxicity response (Fig. 7B). In addition, in many cases we detected transitory drug signatures with high similarity to consensus signatures of different damage responses. Therefore, the concentration-response analysis permits determining the underlying mechanisms of drug toxicity and the concentration ranges where these effects dominate.
3. The off-target signatures of cell damage response were found for most tested drugs, regardless of DILI liability. The critical difference between most- and less-DILI-concern drugs was in the proximity of their plasma concentrations (C_MAX_) to the off-target thresholds (C_OFF_). As a quantitative descriptor, we used the C_OFF_/C_MAX_ ratio termed the ‘safety margin’ (SM). In each class, the most-DILI-concern drugs invariably had the smallest SM values, reliably distinguishing them from less-or no-DILI liability drugs.
4. The TFAP approach helps to explain the rare and unpredictable nature of idiosyncratic DILI. Based on reported C_MAX_ values, the bulk of evaluated most-DILI-concern drugs should not reach the off-target thresholds in vivo, which is consistent with the lack of overt toxicity (Fig. 7). However, the C_MAX_ and C_OFF_ values are not biological constants. The reported C_MAX_ values are average data across the population, whereas individual drug concentrations vary due to multiple environmental and genetic factors. Akin to that, we have approximated the C_OFF_ thresholds in the liver cells based on our data in a model hepatocyte cell line. However, the actual C_OFF_ values can vary among individuals. Because of that, the actual drug concentrations can reach the off-target thresholds and produce adverse effects in some individuals. The probability of that is inversely proportional to the drug safety margin: the smaller the margin, the higher the DILI probability (Fig. 8).
5. Our model agrees with the notion that idiosyncratic DILI stems from a threshold concurrence of drug-specific drivers of toxicity (e.g., drug exposure levels and inherent chemical properties) and individual physiological, environmental, and genetic risk factors(*5, 45*). Furthermore, our mechanistic model is consistent with the phenomenological “rule of three” linking drug propensity for DILI with the dose, lipophilicity, and the formation of reactive metabolites(*10, 46*).
6. In contrast to DILI liability assessments by diverse assay panels for various cell functions, the TFAP approach accounts for multiple damage mechanisms using a single-well assay. Furthermore, the heterogeneous panel data may be difficult to interpret, whereas the TFAP approach produces clear-cut quantitative metrics for DILI liability. Compared to systems biology approaches, e.g., transcriptomics, a distinct advantage of our approach is that it operates with reproducible quantitative signatures that do not require complex bioinformatic analyses and probabilistic predictions.
7. From the practical standpoint, we offer a streamlined decision-making approach for preclinical drug development. The TFAP approach can aid in triaging dangerous candidates and selecting safe drug leads based on their safety margins. The FACTORIAL assay provides clear-cut quantitative estimates for the C_OFF_ values. The other required parameter (C_MAX_) can be obtained from animal studies and *in vitro-in vivo* extrapolation of in vitro assays data(*47*). Furthermore, the TFAP approach can guide lead optimization by assessing chemical modification effects on leads’ safety margins.

**Figure 8.**
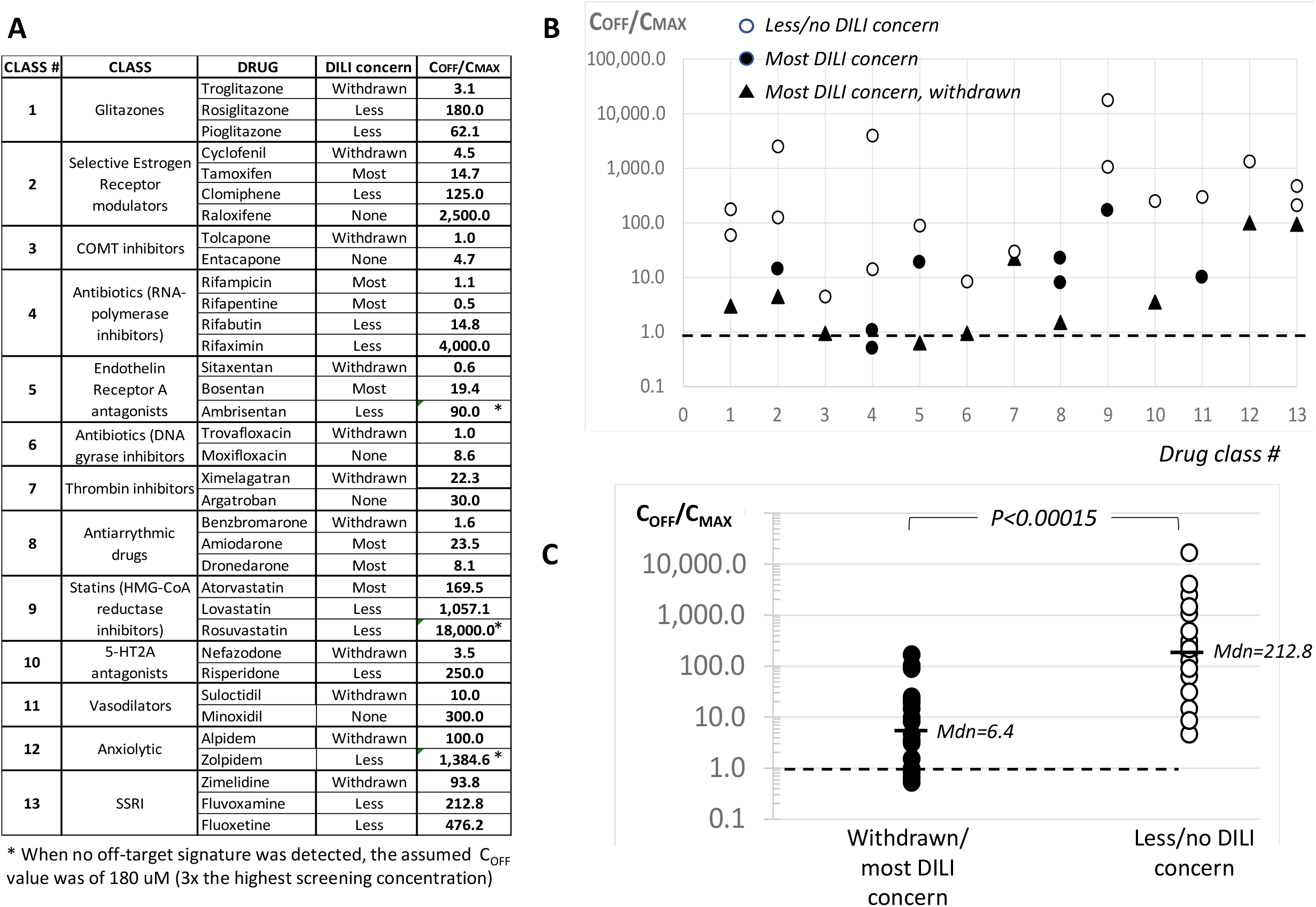
Explaining idiosyncratic drug toxicity by the TFAP approach. The average circulating concentrations (C_MAX_) of most evaluated drugs in the population are below the thresholds of adverse effects (C_OFF_). However, these values can overlap due to the individual variability of C_MAX_ and C_OFF_, causing adverse effects. The probability of that is inversely proportional to the safety margins. Therefore, lower safety margins of most-DILI-concern drugs explain the increased frequency of adverse effects.

**Figure 9.**
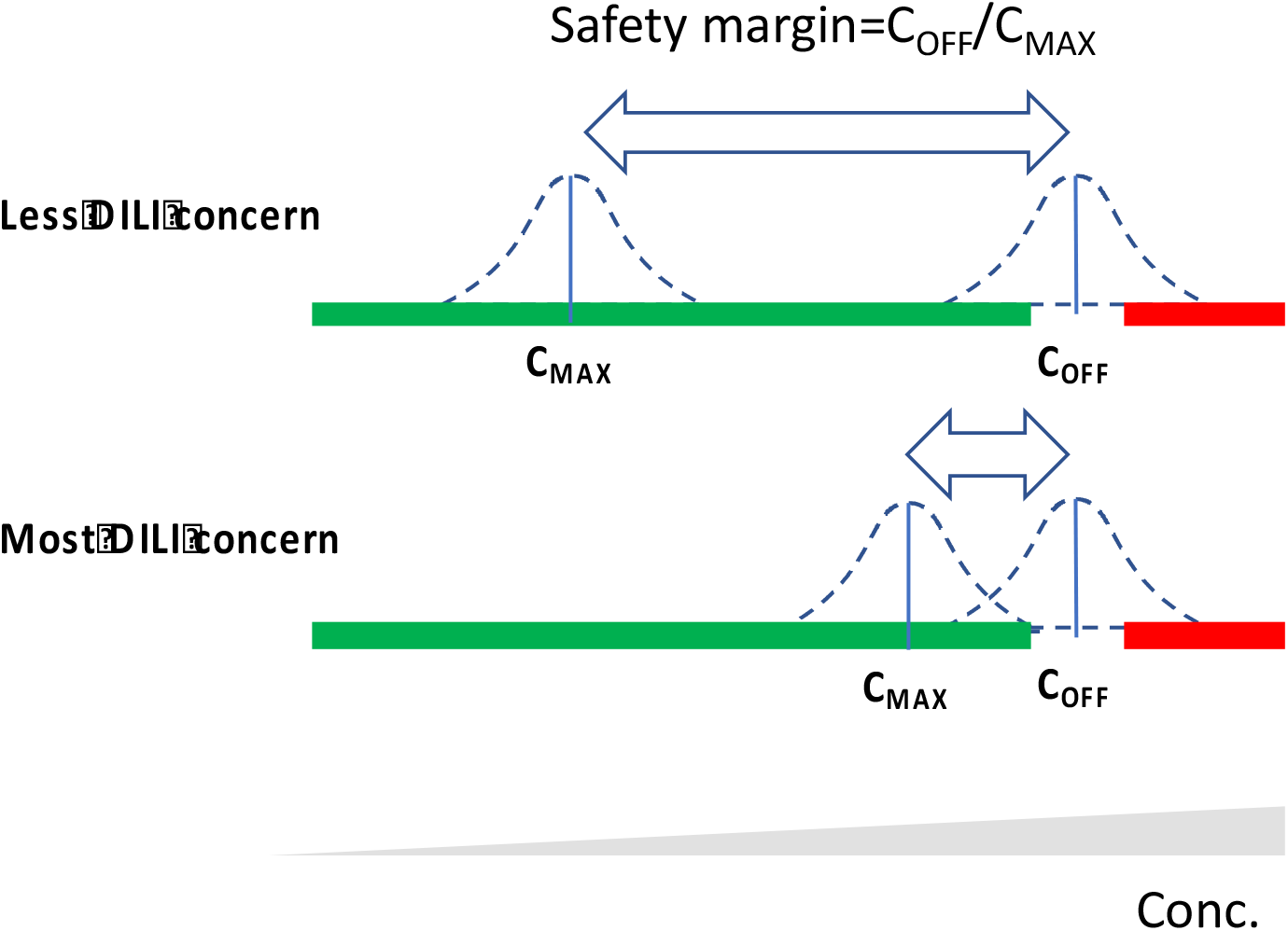

One caveat is that HepG2 cells used in our study have limited metabolic capability(*48*), which could preclude detecting some drug metabolites’ effects. However, there are several ways to obviate this limitation, e.g., by transfecting expression plasmids for drug-metabolizing enzymes(*49*) or using more metabolically active hepatocyte cells(*48*).

In conclusion, the TFAP assessment enables a straightforward DILI liability evaluation approach that complements the existing techniques and streamlines decision-making during the selection and optimization of drug leads.

## MATERIALS AND METHODS

### Cells and reagents

The FACTORIAL assays used the HG19 clone (Attagene) of HepG2 cells (ATCC #HB-8065) selected for an elevated xenobiotic metabolic activity(*14*). Drugs were purchased from Cayman Chemical Company (https://www.caymanchem.com) and Selleck Chemicals (www.selleckchem.com). We used stock solutions of drugs in DMSO, keeping the final DMSO concentration in cell growth medium below 0.2%.

### Cell viability

was evaluated by the XTT (2,3-Bis-(2-Methoxy-4-Nitro-5-Sulfophenyl)-2*H*-Tetrazolium-5-Carboxanilide) assay (ATCC) in HG19/HepG2 cells. As a baseline, we used cells treated with corresponding dilutions of the vehicle (DMSO). The viability data are an average of two replicates. The viability threshold was set at 80% viability.

### The FACTORIAL assay

was conducted as described(*11, 13*). Briefly, the mix of 48 RTU plasmids was transiently transfected using TransIT-LT1 reagent (Mirus) in suspended assay cells, and then the transfected cells were plated into 12-well plates, each well representing an individual FACTORIAL assay. Twenty-four hours later, cells were rinsed and incubated for another 24 h with evaluated compounds in a DMEM growth medium supplemented with 1% charcoal stripped FBS and antibiotics. Total RNA was isolated and analyzed by consecutive steps of RT-PCR amplification, *HpaI* digest, and capillary electrophoresis, as described(*11, 13*) (Fig. S1).

### TFAP signatures

Differential TF activity profiles were calculated by dividing corresponding RTU activity values in compound-treated vs. vehicle-treated cells. The TFAP signatures are radial graphs showing log-transformed fold-changes of RTU activity. Each TFAP signature is an average of three independent FACTORIAL assays.

## Statistical analysis

### Assessing the similarity of TFAP signatures

The pair-wise similarity of TFAP signatures was calculated as the Pearson correlation coefficient *r* (*11, 13*). The signatures with *r* > 0.70 were considered identical, for the probability of two random 48-endpoint signatures having the similarity *r* > 0.70 is very low (*P* < 10^−7^)(*18*). To discriminate the experimental noise, we also calculated the TFAP’s’ Euclidean distance (*d*) from the null signature, considering signatures with d<0.15 as null signatures.

### Calculating the C_OFF_ values

The C_OFF_ threshold for TFAP signature transition was calculated using hierarchical cluster analysis. Specifically, the TFAP signatures of a drug at multiple concentrations were clustered, as previously described(*13*), and the C_OFF_ value was defined as the concentration separating the on-target and off-target clusters.

## Comparing the median safety thresholds across multiple drug classes

The statistical significance (*P-value*) for the median safety margin values across pharmacological classes was calculated by the two-sided Mann-Whitney test using R software.

## Supplementary Data

**Figure S1.**
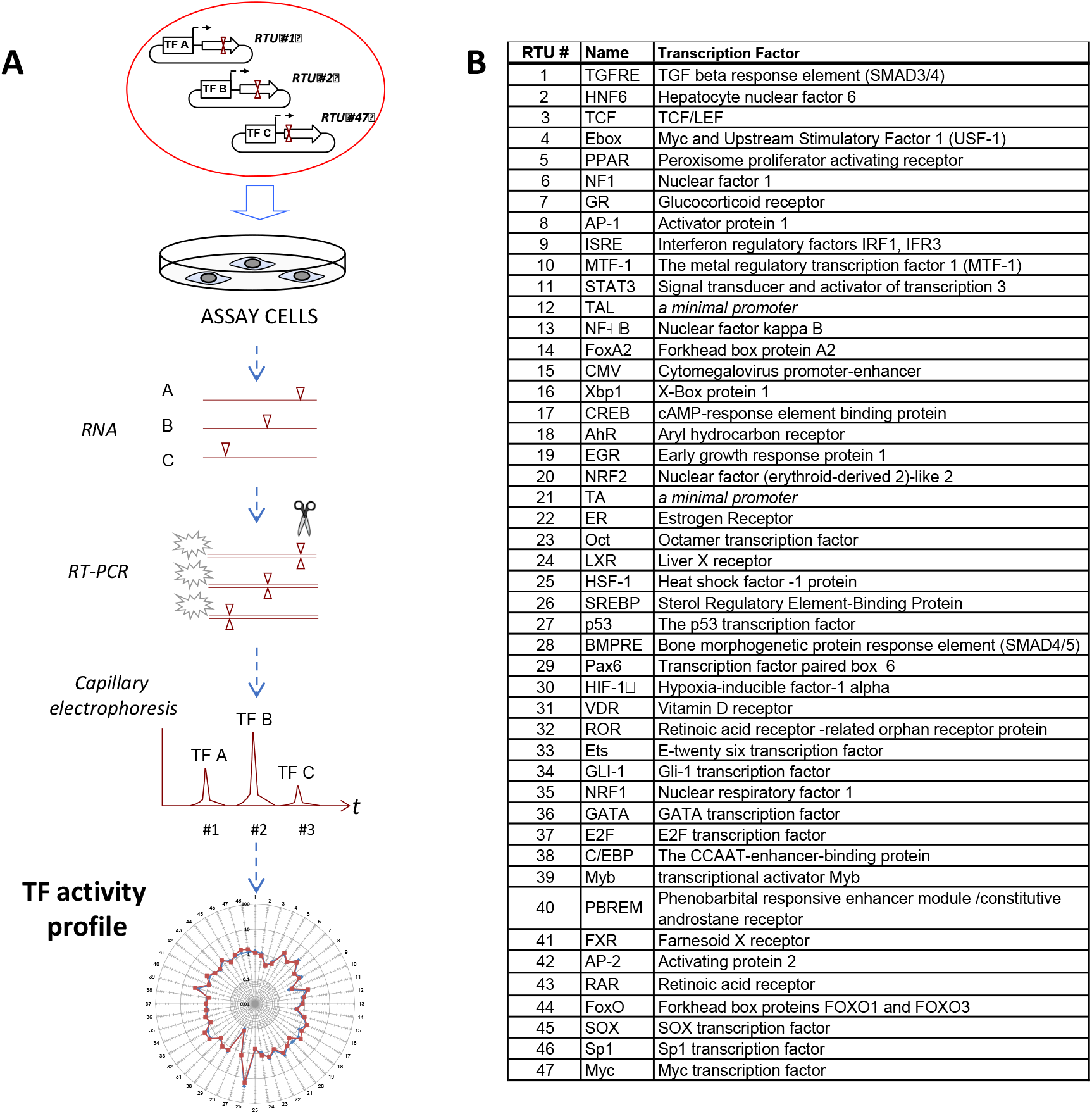

**Table S1.**

| DRUG | C<sub>MAX</sub>,<br>uM | REFERENCES |
| --- | --- | --- |
| Alpidem | 0.2 | PMID 1687121 |
| Ambrisentan | 2.0 | DO - 10.3999/jscpt.41.301 |
| Amiodarone | 0.85 | PMID 18508880 |
| Argatroban | 2.0 | PMID 10730687 |
| Atorvastatin | 0.12 | DOI 10.1093/ndt/gfg048 |
| Benzbromarone | 4.3 | PMID 18508880;21467212 |
| Bosentan | 3.1 | PMID 12047483 |
| Clomiphene | 0.02 | PMID 3607746 |
| Cyclofenil | 0.55 | PMID 17173370 |
| Dronedarone | 0.27 | PMID 21126199 |
| Entacapone | 4.3 | PMID 12588182 |
| Fluoxetine | 0.04 | PMID 16229112 |
| Fluvoxamine | 0.09 | PMID 22931300 |
| Lovastatin | 0.01 | Health Canada Monograph - Lovastatin |
| Minoxidil | 0.2 | PMID 9840212 |
| Moxifloxacin | 7.0 | PMID 22303830;19252339 |
| Nefazodone | 0.9 | PMID 19224530; 15989140;19720731 |
| Pioglitazone | 2.9 | PMID 21702456 |
| Raloxifene | 0.001 | PMID 19224530;19253935 |
| Rifabutin | 0.5 | PMID 7736687 |
| Rifampicin | 9.2 | PMID 21194606 |
| Rifapentine | 14 | DOI 10.1164/rccm.200311-1612OC |
| Rifaximin | 0.005 | https://www.accessdata.fda.gov/drugsatfda_docs/nda/2010/022554Orig1s000ClinPharmR.pdf |
| Risperidone | 0.08 | PMID 19253935 |
| Rosiglitazone | 1.0 | PMID 21467212;19253935 |
| Rosuvastatin | 0.01 | PMID 14693307 |
| Sitaxentan | 31 | PMID 17635499 |
| Suloctidil | 0.22 | DOI 10.1007/978-1-4613-3991-5_10 |
| Tamoxifen | 0.17 | PMID: 22646477; 10649967 |
| Tolcapone | 21 | PMID 15989140 |
| Troglitazone | 6.4 | PMID 15989140;19253935 |
| Trovafloxacin | 7 | PMID 15989140 |
| Ximelagatran | 0.3 | PMID 17532684 |
| Zimelidine | 0.6 | PMID:6228368 |
| Zolpidem | 0.13 | PMID 12919178 |

**Figure S2.**
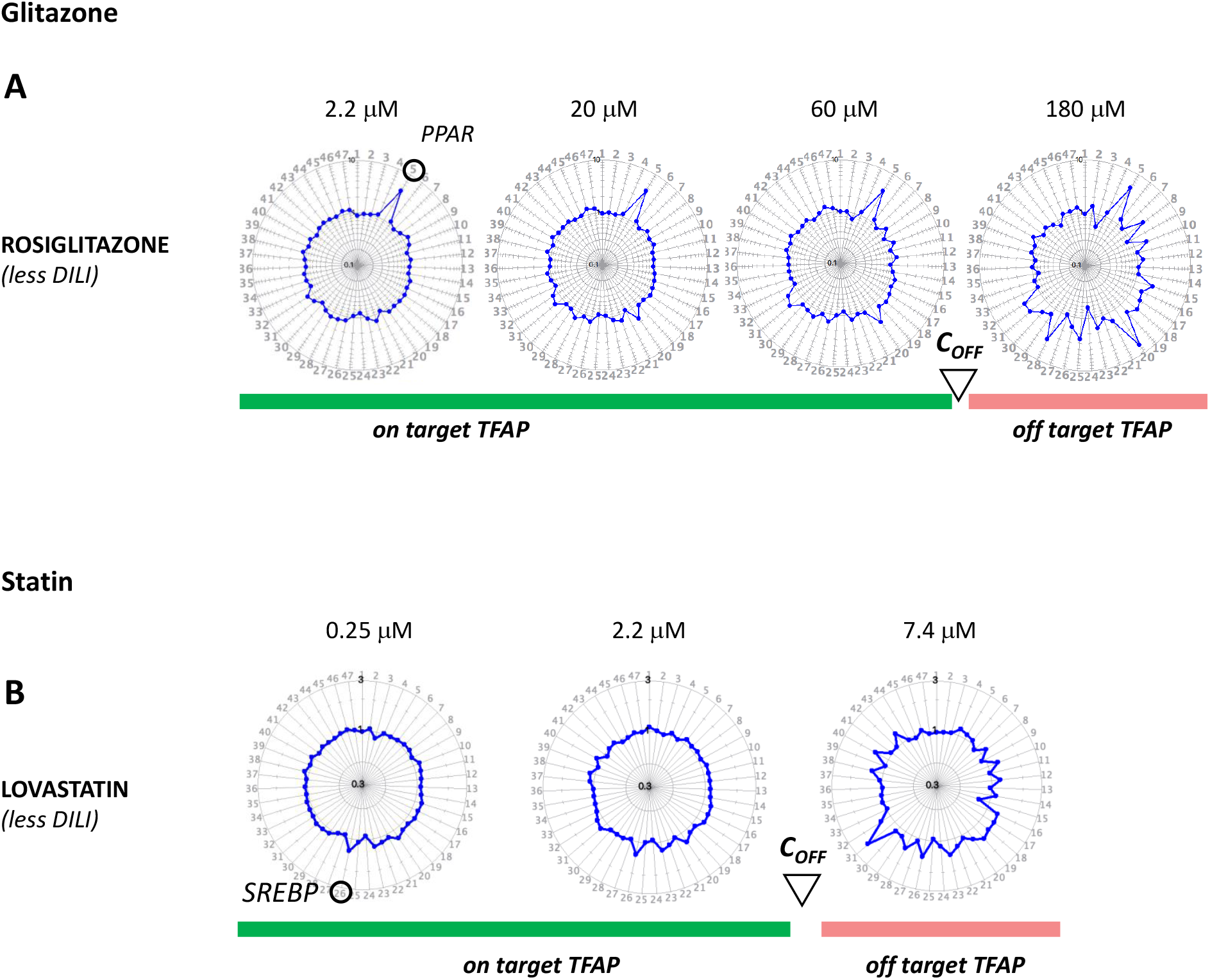
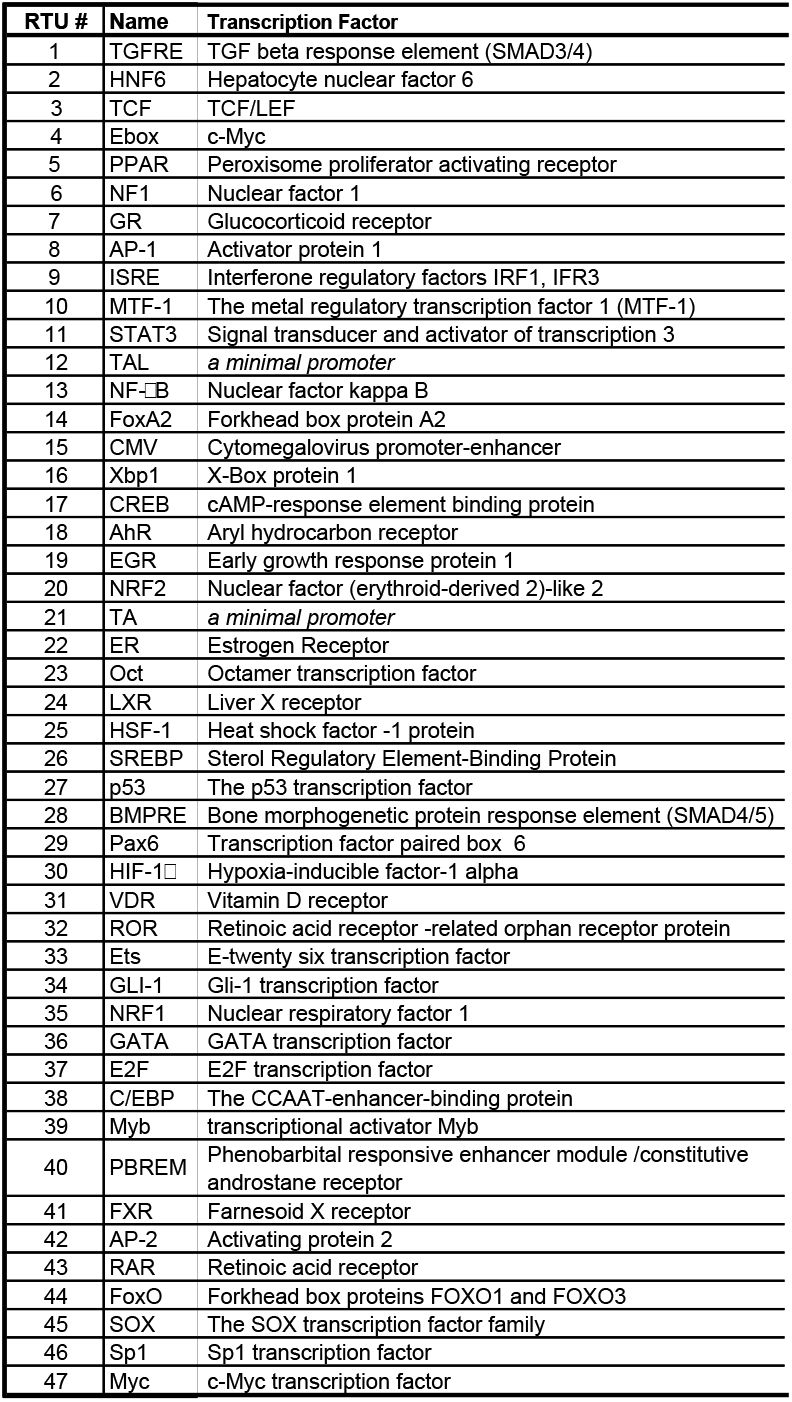

**Figure S3.**
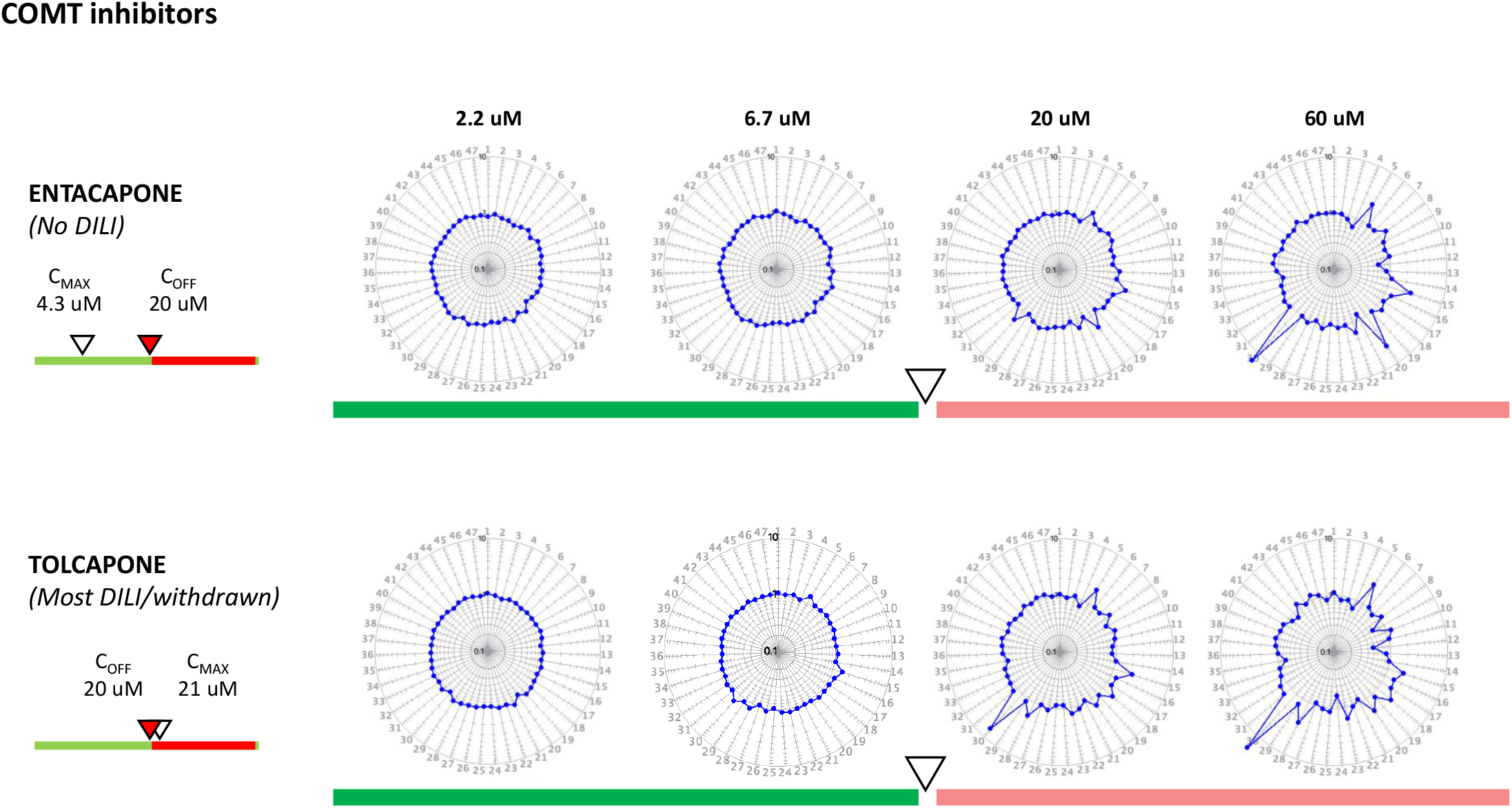

**Figure S4.**
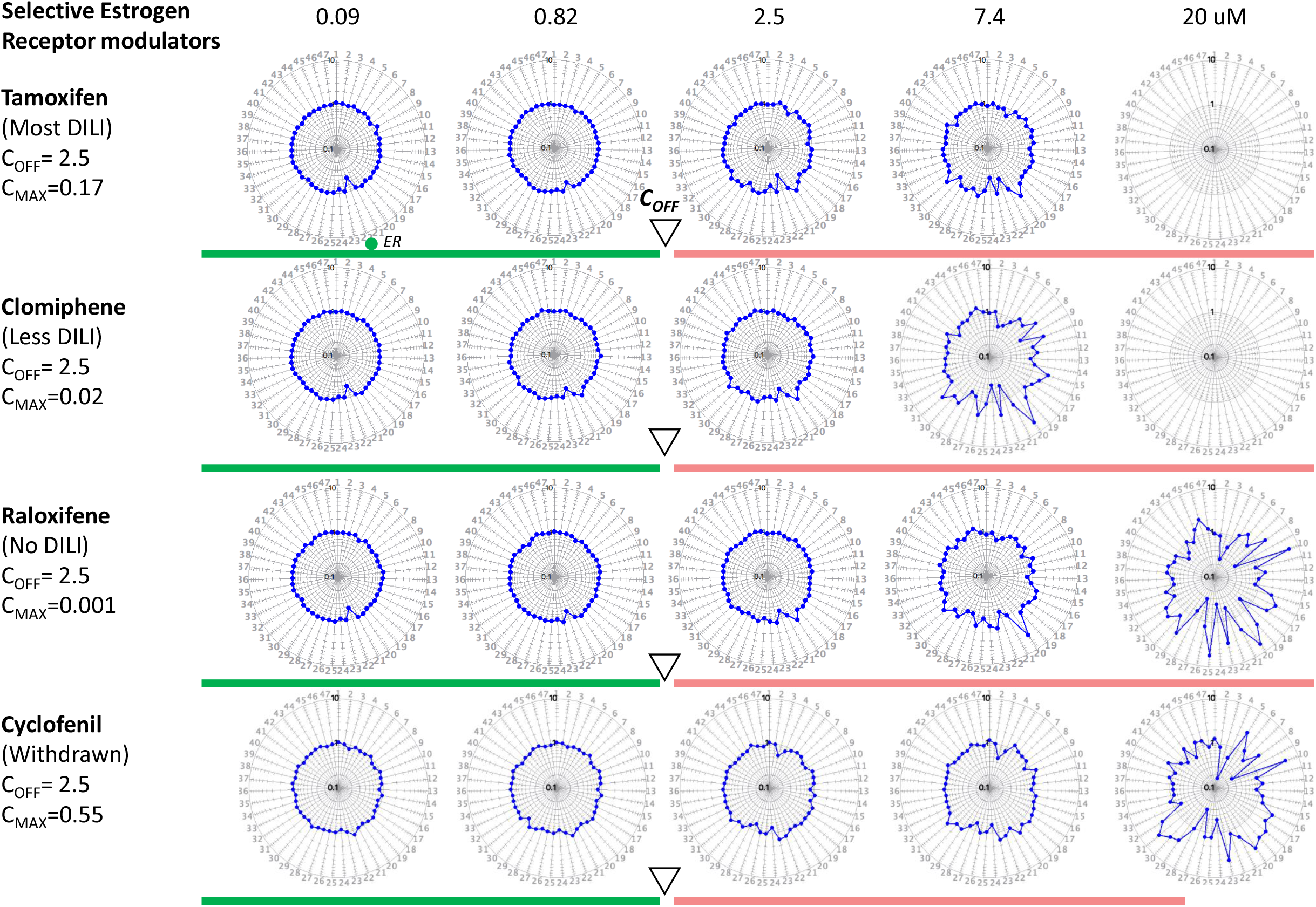

**Figure S5.**
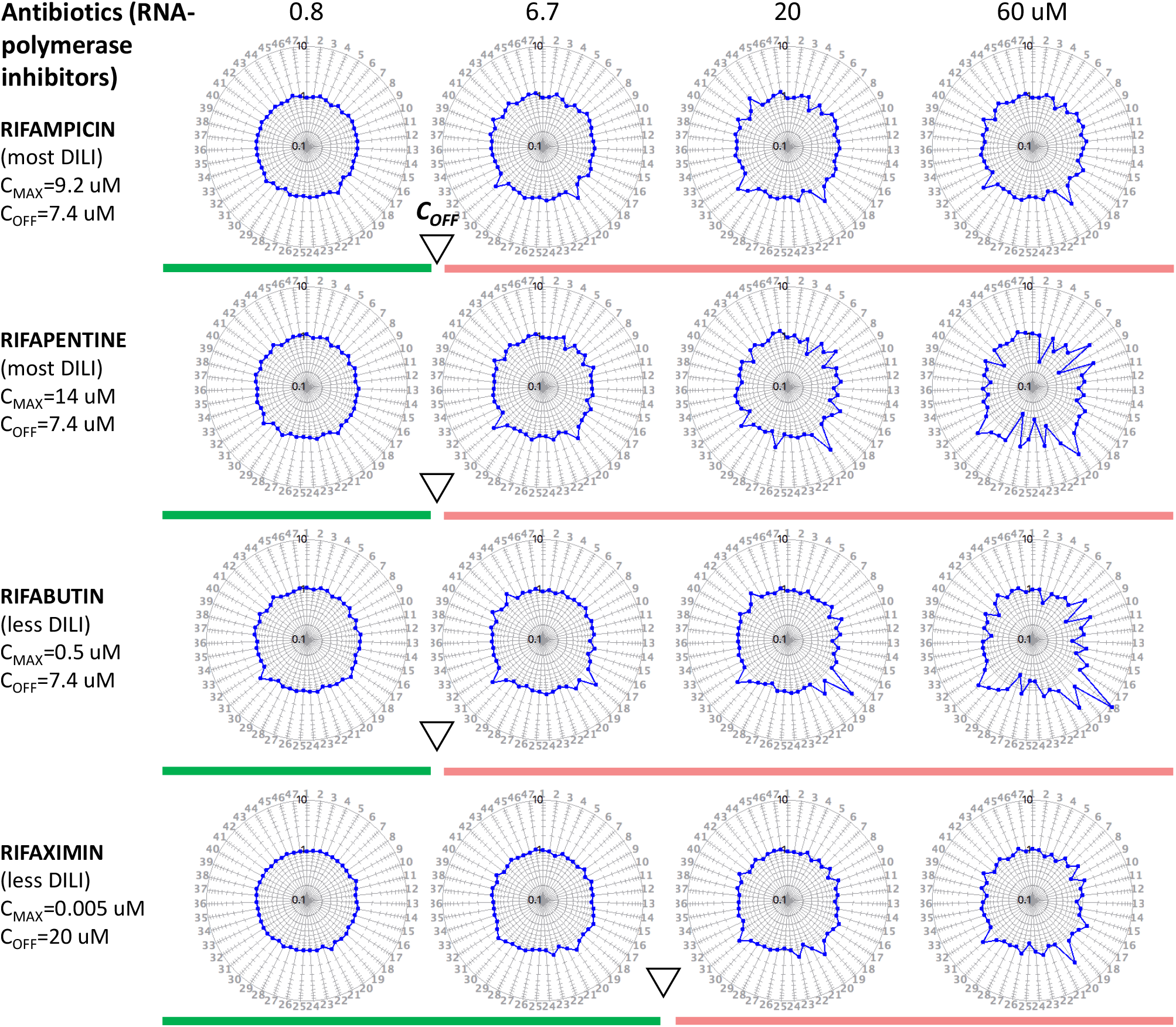

**Figure S6.**
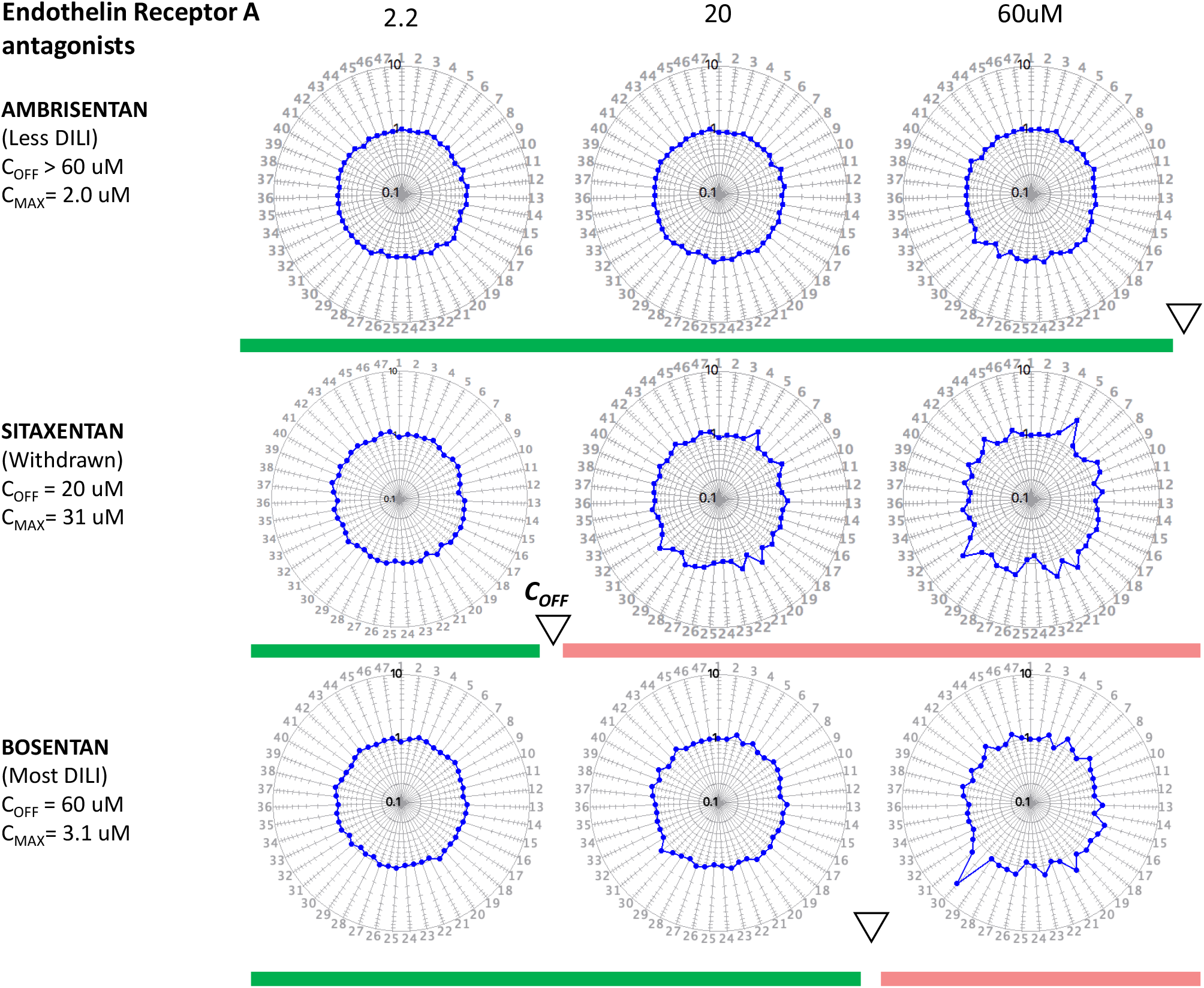

**Figure S7.**
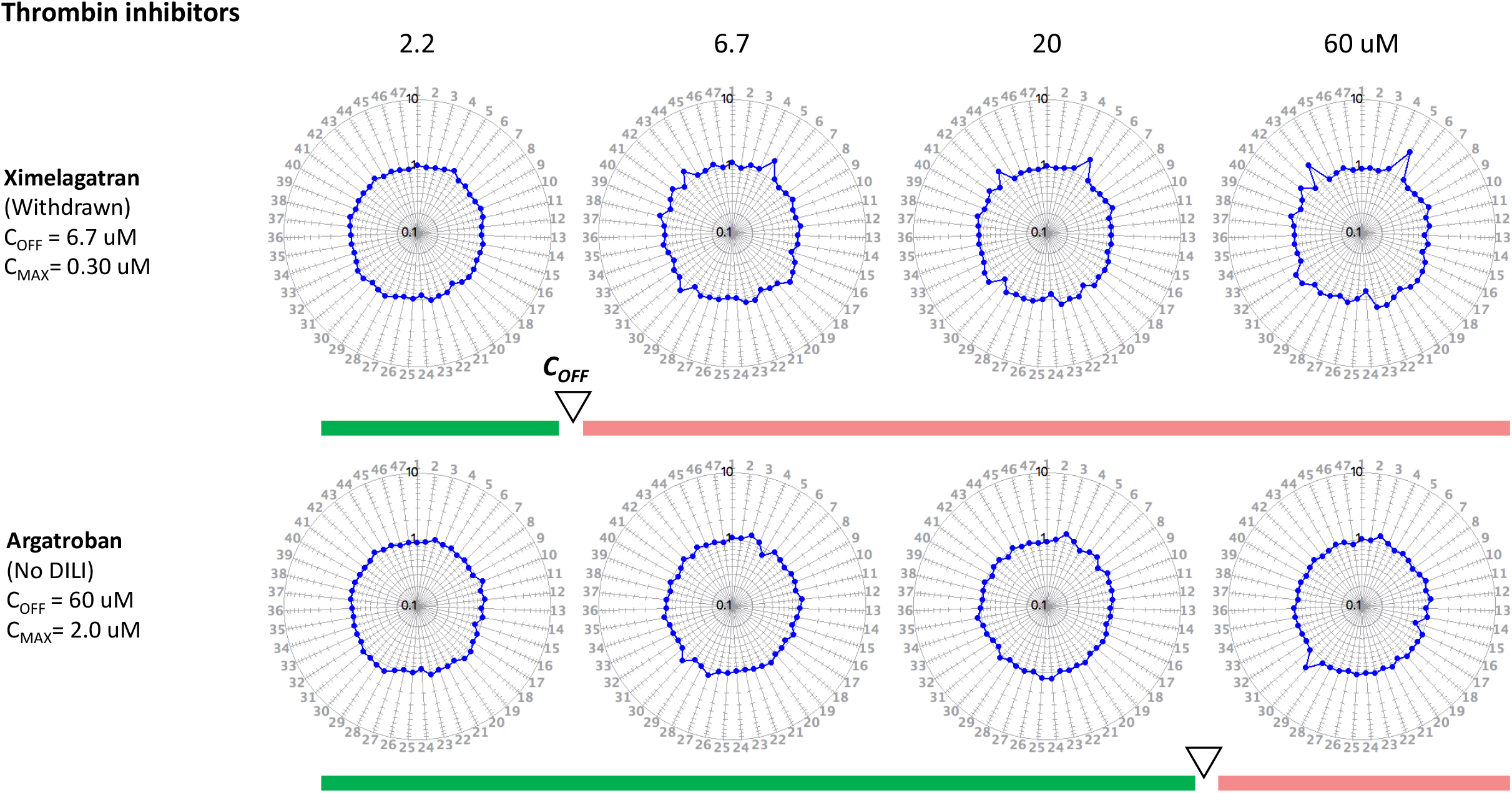

**Figure S8.**
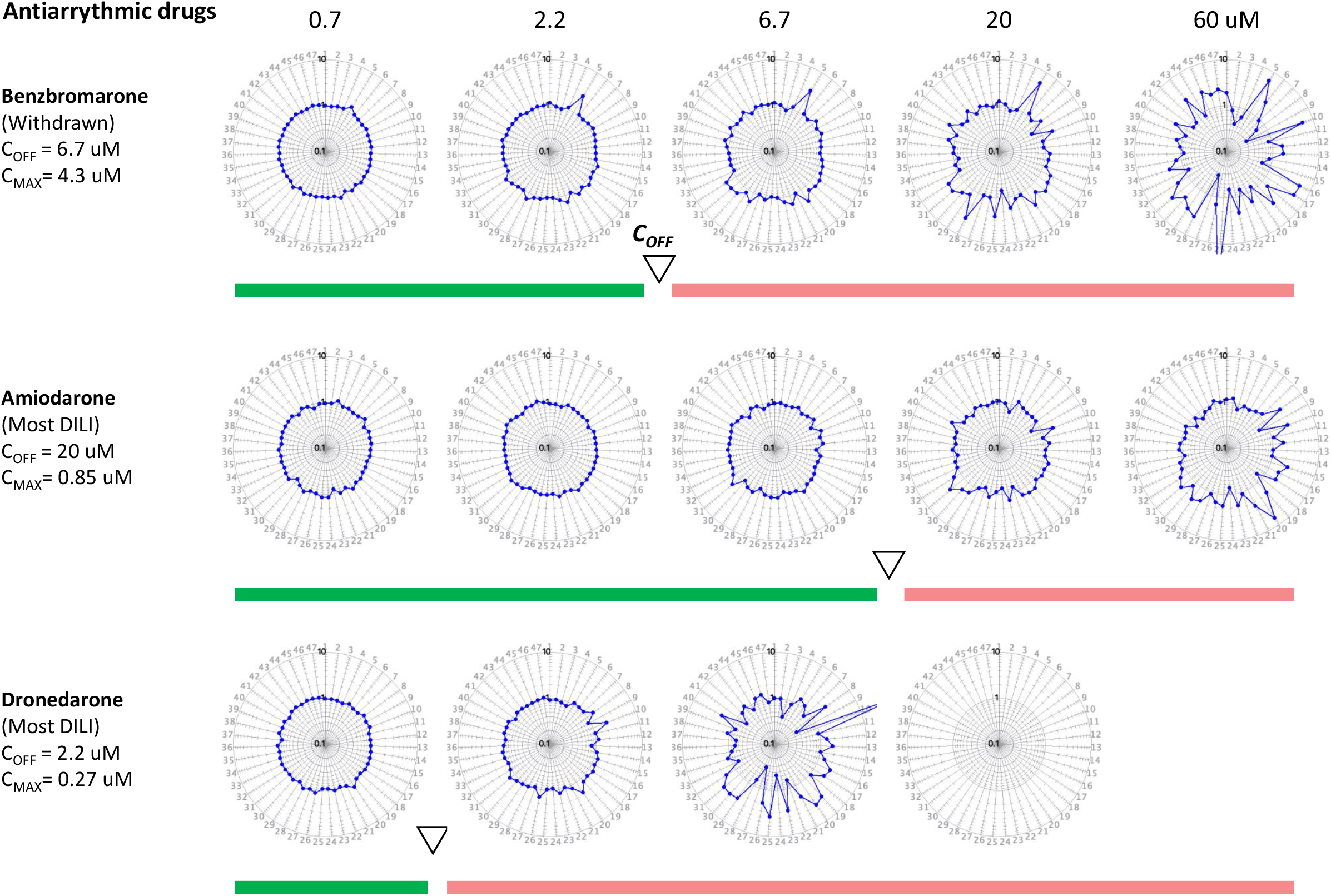

**Figure S9.**
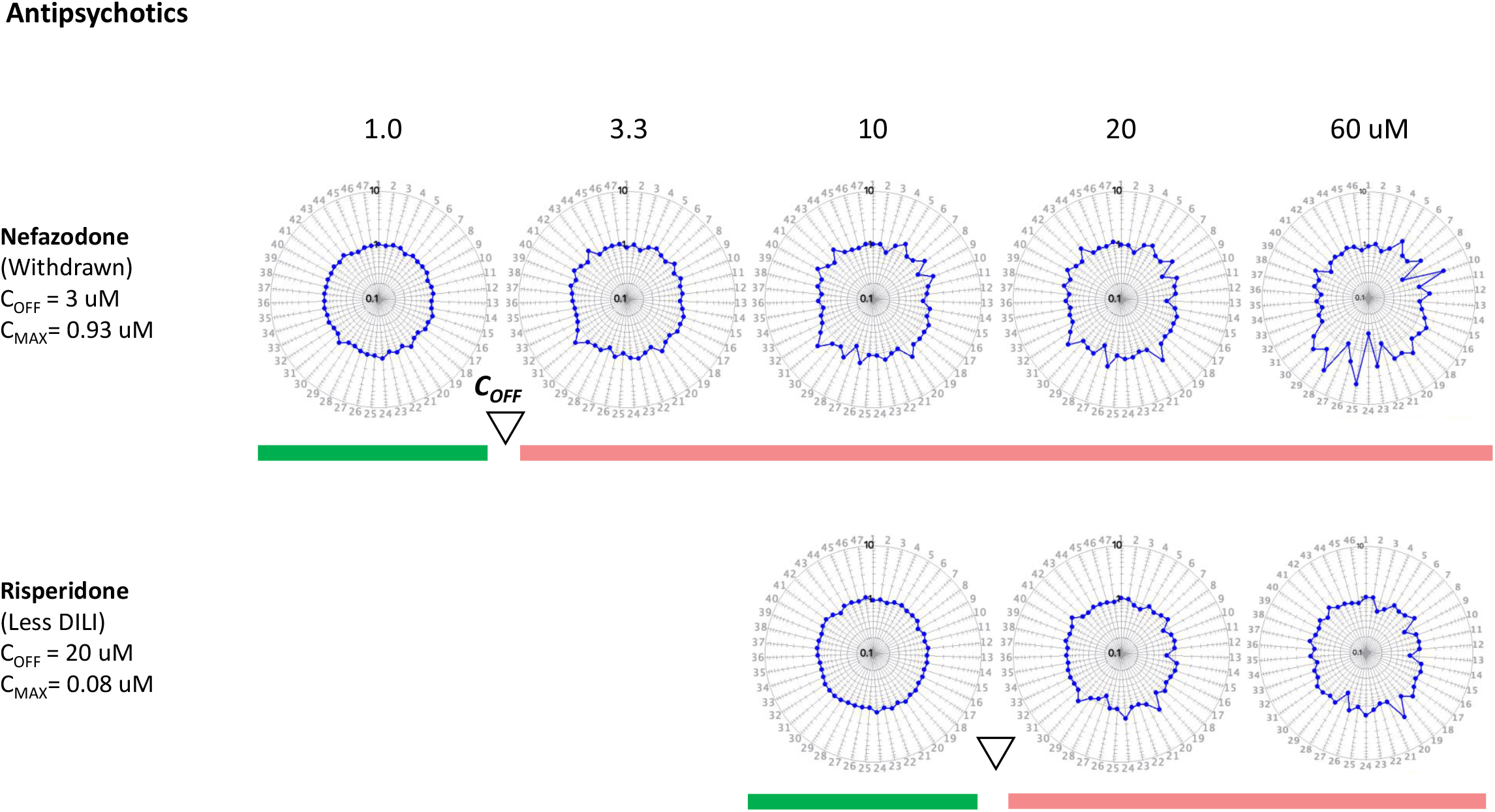

**Figure S10.**
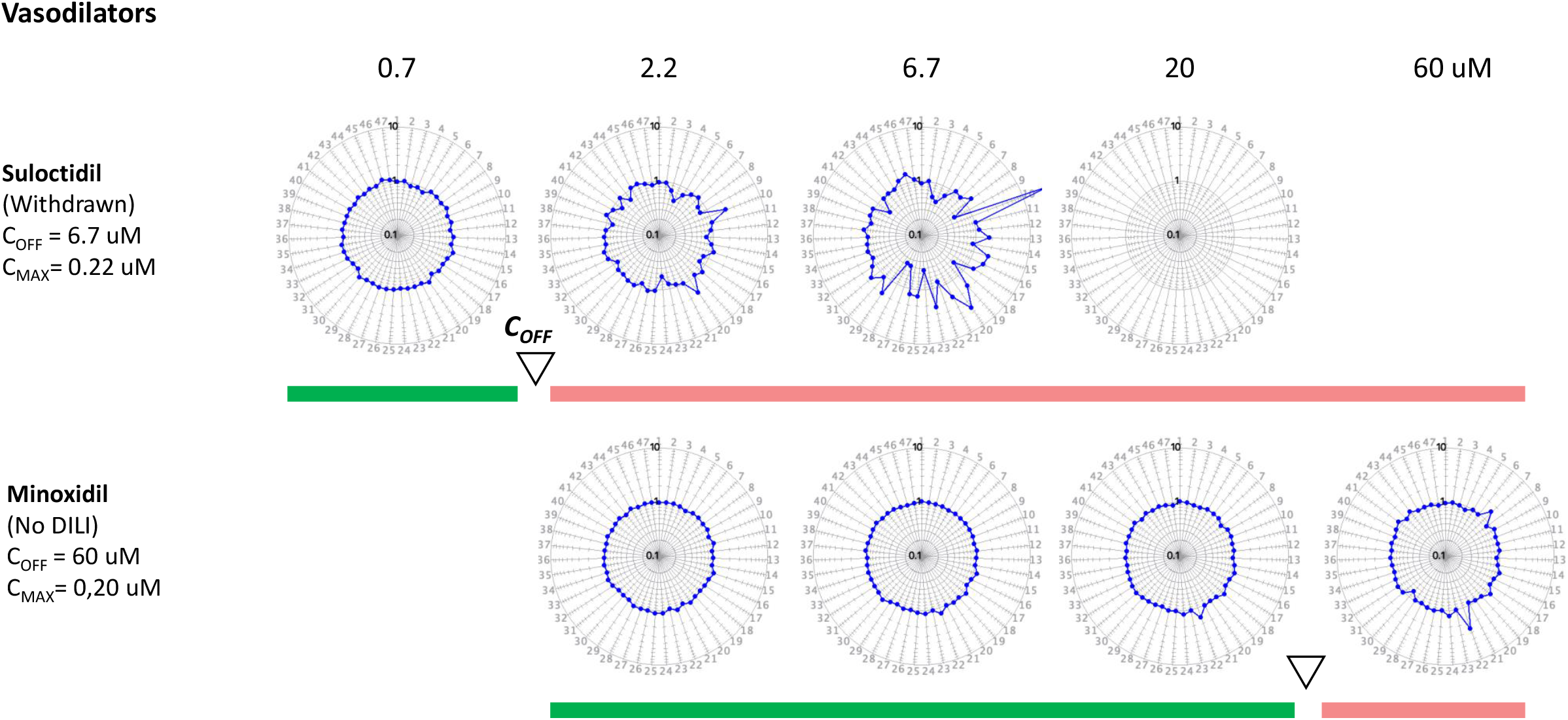

**Figure S11.**
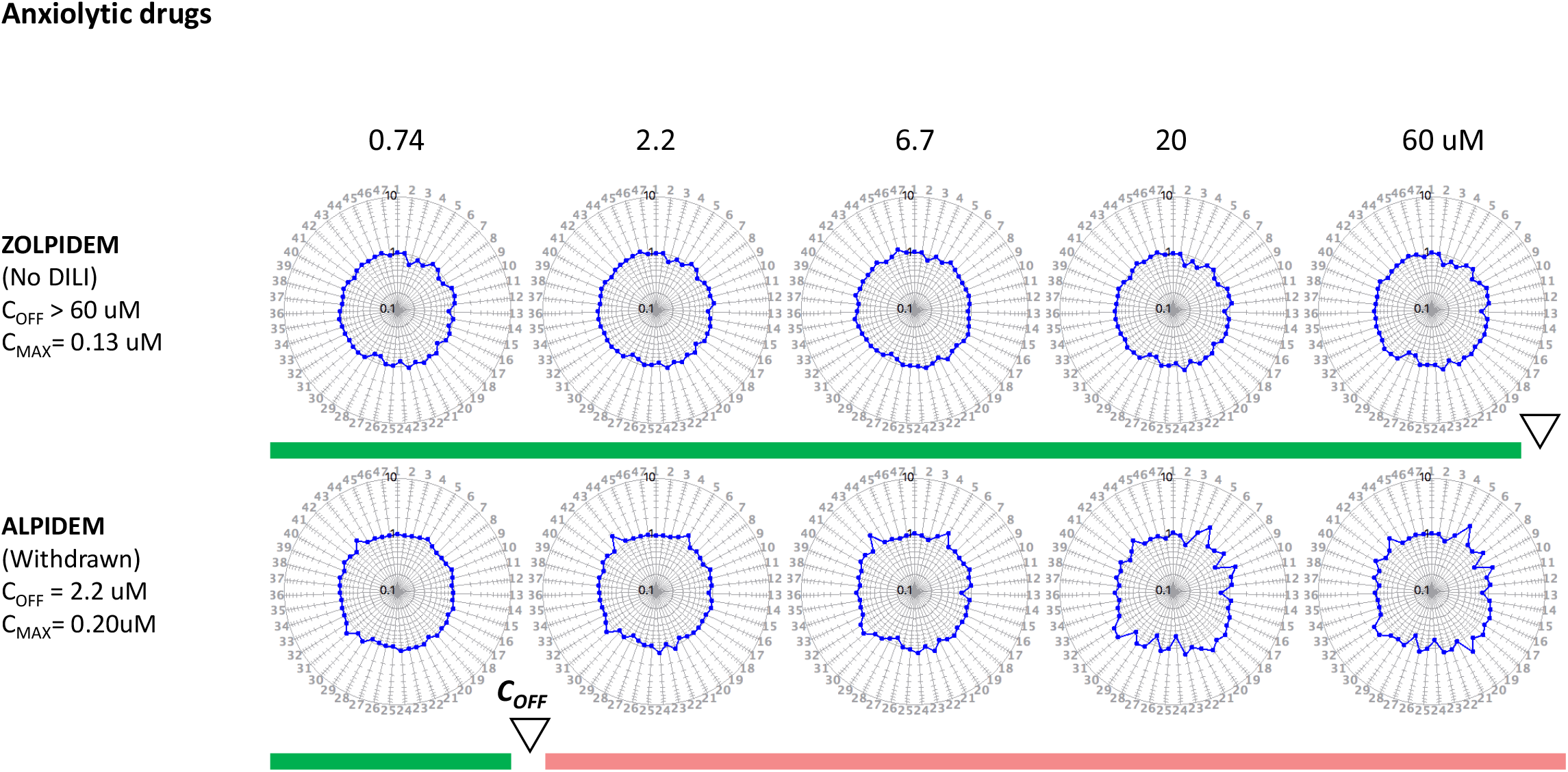

**Figure S12.**
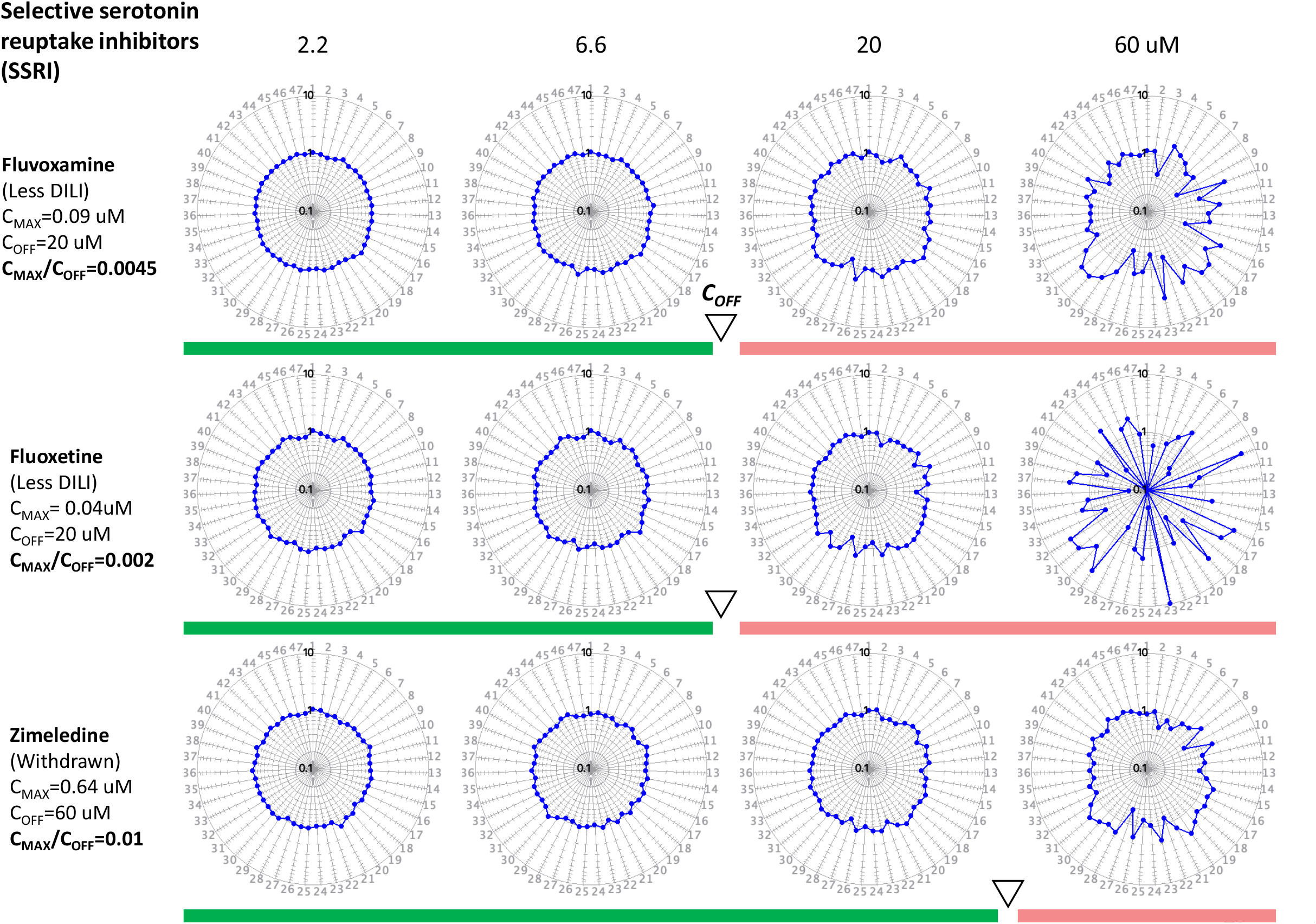

